# Development and Validation of the Mentoring in Undergraduate Research Survey

**DOI:** 10.1101/2023.08.19.553952

**Authors:** Lisa B. Limeri, Nathan T. Carter, Riley A. Hess, Trevor T. Tuma, Isabelle Koscik, Alexander J. Morrison, Briana Outlaw, Kathren Sage Royston, Benjamin H. T. Bridges, Erin L. Dolan

## Abstract

Here we present the development and initial validation of the Mentoring in Undergraduate Research Survey (MURS) as a measure of a range of mentoring experienced by undergraduate science researchers. We drafted items based on qualitative research and refined the items through cognitive interviews and expert sorting. We used national dataset to evaluate the internal structure of the measure and a second national dataset to examine how responses on the MURS related to theoretically-relevant constructs and student characteristics. Our factor analytic results indicate seven lower order forms of mentoring experiences: abusive supervision, accessibility, career and technical support, psychosocial support, interpersonal mismatch, sexual harassment, and unfair treatment. These forms of mentoring mapped onto two higher-order factors: supportive and destructive mentoring experiences. Although most undergraduates reported experiencing supportive mentoring, some reported experiencing absence of supportive as well as destructive experiences. Undergraduates who experienced less supportive and more destructive mentoring also experienced lower scientific integration and a dampening of their beliefs about the value of research. The MURS should be useful for investigating the effects of mentoring experienced by undergraduate researchers and for testing interventions aimed at fostering supportive experiences and reducing or preventing destructive experiences and their impacts.

**Highlight summary:** This study presents the development and initial validation of the Mentoring in Undergraduate Research Survey, including evidence of its internal structure as well as convergent, discriminant, and predictive validity.

## INTRODUCTION

Proponents of undergraduate STEM education reform advocate for widespread involvement of undergraduates in research because of the potential for professional, academic, and personal benefits (AAAS, 2011; Byars-Winston et al., 2015; Estrada et al., 2018). Undergraduate research experiences (UREs) are also increasingly recognized for their capacity to promote the integration of students into the scientific community, especially students from marginalized or minoritized backgrounds (Estrada et al., 2011; Hernandez et al., 2017b; Hernandez, Woodcock, et al., 2018). Multiple qualitative and quantitative studies have shown that mentoring plays a critical role in STEM undergraduate researchers’ personal and professional development (Aikens et al., 2016; Estrada et al., 2018; Hernandez, Hopkins, et al., 2018; Joshi et al., 2019; Thiry & Laursen, 2011). Studies examining mentoring in UREs have generally focused on positive mentoring that undergraduates experience (Aikens et al., 2016, 2017; Hernandez et al., 2017b). However, studies from both workplace and academic settings have shown that not all mentoring experiences are positive.

Mentoring, like any interpersonal relationship, can include dysfunctional elements or problematic events, which are collectively referred to as negative mentoring (Eby et al., 2000; Kram, 1983; Scandura, 1998; Simon & Eby, 2003). Workplace mentees report problems with mentors such as personality mismatches, mentor neglect, and mentor sabotage, as well as mentors lacking expertise (Eby et al., 2000, 2004; Simon & Eby, 2003). Doctoral students in the life sciences also report negative mentoring experiences with their research advisors, including inaccessibility, deceit, and problematic supervisory styles such as micromanagement (Tuma et al., 2021). A few studies of mentoring in UREs have noted variation in the quality of mentoring, such as absenteeism, unrealistic expectations, or insufficient guidance from mentors (Bernier et al., 2005; E. L. Dolan & Johnson, 2010; Thiry & Laursen, 2011). To define negative mentoring in undergraduate research, we previously conducted a qualitative study of negative mentoring experiences among undergraduate researchers in the life sciences (Limeri, Asif, Bridges, et al., 2019). We identified seven types of negative mentoring experiences: absenteeism, abuse of power, interpersonal mismatch, lack of career support, lack of psychosocial support, misaligned expectations, and unequal treatment. Although this research characterized negative mentoring experiences among undergraduate researchers, it did not provide direct evidence of the effects of these experiences.

Studies from the workplace suggest that negative mentoring harms mentees, decreasing their job satisfaction and increasing their stress as well as their intentions to leave their jobs (Eby & Allen, 2002). One study indicated that workplace negative mentoring may be so damaging that mentees who experience it may be worse off than if they had no mentor at all (Ragins et al., 2000). Negative mentoring is most strongly associated with negative mentee outcomes when the mentoring relationships are assigned rather than formed organically (Eby & Allen, 2002). This is concerning because formal assignment is often how mentoring relationships are formed in UREs; either a faculty member assigns an undergraduate to a graduate or postdoctoral mentor or an undergraduate is assigned to a faculty member’s research group (Dolan & Johnson, 2009; Erickson et al., 2022; Limeri, Asif, & Dolan, 2019).

Quality mentorship during UREs is especially beneficial for minoritized students (Estrada et al., 2018; Hernandez et al., 2017a). These findings raise concerns that negative mentoring experiences could disproportionately affect minoritized students and exacerbate inequities in STEM. Research indicates that UREs are especially beneficial for students from minoritized or marginalized backgrounds because these experiences promote a sense of fit with the scientific community (Estrada et al., 2011, 2018; Hurtado et al., 2009). Negative mentoring experiences may prevent rather than promote a sense of belonging, and thus disproportionately harm students already facing barriers to their integration in STEM.

### Measuring Negative Mentoring

Given the widespread recommendations to involve undergraduate STEM students in research and the potential for negative mentoring to cause harm (Gentile et al., 2017), understanding the range of mentoring that undergraduate researchers experience and how it affects them is critical. Accomplishing this requires a measure of the range of mentoring experienced by undergraduate researchers that is supported by strong evidence of validity. Here we report the development of and initial construct validity for such a measure: the <u>M</u>entoring in <u>U</u>ndergraduate <u>R</u>esearch <u>S</u>urvey (MURS). We opted to develop a new measure because we were unable to identify existing instruments suitable for measuring the range of mentoring experienced by undergraduate researchers that we observed in our qualitative work (Limeri, Asif, Bridges, et al., 2019).

Several instruments have been used to measure mentorship quality (reviewed in Byars-Winston & Dahlberg, 2019); yet the majority lack evidence of validity and reliability (Hernandez, 2018). Few if any have been designed or used to assess mentorship quality at the undergraduate level. Furthermore, most have been designed to assess positive mentorship, and thus are likely to fall short of capturing key elements of negative mentoring experiences. For example, the Mentoring Competency Assessment was developed to evaluate research mentors’ skills before and after a mentoring training program (Fleming, House, Hanson, et al., 2013). The scale asks mentees to rate their mentor’s ability on 26 skills associated with six mentor competencies: maintaining effective communication, aligning expectations, assessing understanding, addressing diversity, fostering independence, and promoting professional development. These competencies align with some but not all of the negative mentoring experienced by undergraduate researchers in our prior work. For instance, we found that students reported mismatches with their mentor’s personality or work style (Limeri, Asif, Bridges, et al., 2019), which are unrelated to mentor skills per se. In addition, undergraduates most often reported mentor absenteeism as a form of negative mentoring, which mentees often attributed to mentors being overcommitted rather than unskilled. In sum, there does not appear to be a measure suitable for investigating how negative mentoring experiences affect undergraduate students or the outcomes they realize from participating in UREs.

### Measurement Validity Framework

Here we report the development and evidence of validity for the MURS as a measure of mentoring experiences for use with undergraduate science researchers. To guide the development process, we adopted Kane’s argument-based approach to measurement validity (Kane et al., 1999). In this framework, validity is not an inherent property of a measurement instrument, but rather an argument for a proposed interpretation of responses to an instrument, which must be supported by evidence. The interpretive argument in this case is that undergraduate researchers’ responses to the items on the MURS are indicative of students’ mentoring experiences, such that students with higher scores experienced more of that form of mentoring. The process of building the validity argument involves identifying and providing evidence in support of the assumptions underlying this argument. Here we provide evidence to support this and other assumptions to build a validity argument for the MURS.

## METHODS AND RESULTS

Studies of mentoring experiences as well as related experiences of abusive supervision and workplace incivility have primarily operationalized these phenomena in terms of recipients’ perceptions (Eby et al., 2013; Schilpzand et al., 2016; Tepper et al., 2017). Although perceptions have been criticized for their lack of objectivity (Linn et al., 2015; Tepper, 2000), we have chosen to use this same approach here for multiple reasons. First, directly observing mentoring would be intrusive and impractical, and negative mentoring may not always be visible to observers. Second, mentors may not be aware that particular behaviors are problematic and may not be willing to report less-than-ideal behavior, making mentor reports of negative mentoring equally subjective. Finally, mentee perceptions of mentoring have been shown to fundamentally alter these relationships and to have long-term effects on mentee outcomes (Eby et al., 2008, 2010; Eby & Allen, 2002; Scandura, 1998). Thus, our intent is to measure undergraduate researchers’ perceptions of their mentoring experiences.

We carried out the process of developing and collecting validity evidence for the MURS over three phases: substantive, structural, and external (Benson, 1998). All phases of the study were reviewed and determined to be exempt by the University of Georgia Institutional Review Board (STUDY00004954). For ease of reading, we present the methods and results together for each phase. In the final phase, we also begin to investigate how mentoring experiences influence undergraduate researchers’ integration into the scientific community.

### Substantive Phase

Our aim with this phase was to collect evidence of content-related and response process validity for the MURS (Messick, 1995). We started by defining and characterizing negative mentoring by identifying observations that reflect the construct (Benson, 1998). Specifically, we carried out a qualitative characterization of negative mentoring experienced by undergraduate researchers to define the content domain of the construct (Limeri, Asif, Bridges, et al., 2019). This work was a useful foundation for capturing a range of mentoring experiences because undergraduates reported both the absence of supportive experiences and mentor behaviors, characteristics, or interactions they experienced as actively harmful or destructive. For comprehensibility and ease of comparison, we present our methods and results using a common, negative valence such that supportive experiences are described in terms of their absence and destructive experiences are described in terms of their presence.

We drafted 107 survey items that corresponded to the seven dimensions of mentoring experiences identified in Limeri et al. (2019): absenteeism, which we renamed inaccessibility (13 items), abuse of power, which we renamed abusive supervision (25 items), interpersonal mismatch (13 items), insufficient career and technical support (12 items), insufficient psychosocial support (14 items), misaligned expectations (18 items), and unequal treatment (12 items). We also adapted five items to represent an eighth dimension, sexual harassment, resulting in 112 items altogether. The sexual harassment items were preceded with a content warning and based on items previously used to measure undergraduates’ experiences with sexual harassment in academic settings (Aycock et al., 2019).

We pilot tested the 112 items by conducting cognitive interviews with undergraduate researchers, which provided evidence of response process validity (i.e., the items were understood as intended). Using a screening survey (see Supplemental Materials), we recruited 32 participants from 14 institutions who had experienced a range of mentoring quality, from mostly positive to mostly negative. Of these, we selected 15 participants from a diverse group of 11 institutions: 6 very high research activity, 1 high research activity, 2 master’s-granting institutions, 1 community college, and 1 research institute; 3 of the institutions were classified as minority-serving (Indiana University Center for Postsecondary Research, n.d.). Participants were compensated with a $25 gift card. Because of the large number of items, each participant reviewed only a subset (one or two dimensions for a total of 15-25 items) such that each item was reviewed by three or four participants. Based on the cognitive interviews (questions provided in Supplemental Materials), we refined and revised the items. We ultimately selected 57 items that were most clearly and consistently interpreted and best represented the range of the construct.

As a final step in the substantive phase, we conducted a Q-sort activity (Nahm et al., 2002) with nine individuals to provide further evidence of content validity. These individuals were selected based on their expertise in mentoring research or extensive experience mentoring undergraduate researchers. After randomly sorting the items and providing definitions for each dimension, we asked the experts to assign each of the 57 items into one of the eight dimensions. We set 70% agreement among the experts as a threshold for retaining the item as is; 40 items passed this threshold. We also asked the experts to indicate their confidence in their assignment and to offer their expert judgment of the relevance of the item to the assigned dimension. For the 40 items with high agreement, the associated certainty and relevance ratings were high (i.e., 70% threshold for ratings of “high” relevance and certainty was reached for all of these items), indicating that we were capturing the main ideas underpinning each dimension. We reworded the remaining 17 items to address ambiguities, producing 57 items that reflected the eight dimensions.

### Structural Phase

Our aim with this phase was to begin to produce evidence of the construct-related validity of the MURS (Messick, 1995). To accomplish this, we examined the extent to which the observed variables (i.e., item responses) covaried among themselves and compared that structure to the theorized seven dimensions of undergraduate mentoring experiences and sexual harassment (Benson, 1998).

We also collected personality data based on the Big Five model of personality traits, namely openness, conscientiousness, extraversion, agreeableness, and neuroticism, which is the dominant model of personality structure in psychological research (John, 2021). We reasoned that undergraduates might vary in their perceptions or reporting of experiences or in their interactions with mentors as a result of their personality traits. For instance, the trait of neuroticism includes the tendency toward negative feelings. Individuals high on neuroticism can interpret ordinary situations as threatening (Widiger & Oltmanns, 2017) and show heightened sensitivity to social cues and relationship conflict (Denissen & Penke, 2008). Thus, undergraduates with elevated levels of neuroticism may experience interactions with mentors more negatively. Individuals high on conscientiousness, or the tendency to be diligent and take obligations seriously, tend to be perceived by supervisors and peers as more engaged in their jobs and their work is more highly rated (Bakker et al., 2012). Thus, undergraduates high on conscientiousness may garner more accolades and support from mentors, reducing their likelihood of reporting negative mentoring experiences. By analyzing relationships between responses on the MURS and personality traits, we sought to explore the possibility that the MURS was measuring facets of personality that might make individuals more or less likely to report negative experiences with mentors.

#### Recruitment and Data Collection

We recruited by email a national sample of undergraduates who had indicated they had completed at least one term (quarter, semester, summer) of mentored research within the past year (Table 1) to respond to the 57 MURS items. We received 573 survey responses in total, of which 16 did not consent to be included in the study and thus were removed from the analysis. We included attention checks (items that directed respondents to select a particular response, e.g., “This is a control question, please select ‘strongly agree’”) to screen out responses that reflected insufficient attention (DeSimone et al., 2015). Respondents had to select both attention check responses accurately to be included in the analysis; 36 responses were excluded because they did not pass one or both attention checks. Thus, the final analytic sample was n=521. Students took an average of 15 minutes to complete the survey and were compensated with a $10 gift card. Our survey included our 57 MURS items as well as a 20-item measure of the Five-Factor model of personality (mini-IPIP; Donnellan et al., 2006) and a series of demographic questions (see Supplemental Materials).

**Table 1.** Demographic information for participants surveyed in the structural and external phases. Institution type was determined using the Carnegie Classification of Institutions (Indiana University Center for Postsecondary Research, n.d.). Racial/ethnic identity counts may not add up to 100% because participants could select multiple racial/ethnic identities.

| **Demographic** | **Structural Phase Participants**<br>**(n=521)** | **External Phase Participants**<br>**(n=348)** |
| --- | --- | --- |
| *Institution Type* | *Reported by 518 (99%) at 60 institutions* | *Reported by 348 (100%) at 32 institutions* |
| Very High Research Activity | 347 (67%) at 22 institutions | 284 (82%) at 22 institutions |
| High Research Activity | 57 (9%) at 6 institutions | 53 (15%) at 4 institutions |
| Doctoral Universities | 0 | 1 (0.3%) at 1 institution |
| Masters-Granting | 30 (5.8%) at 9 institutions | 7 (2.0%) at 2 institutions |
| Primarily Undergraduate | 60 (12%) at 12 institutions | 3 (0.9%) at 3 institutions |
| Community College | 24 (4.6%) at 11 institutions | 0 |
| *Race/Ethnicity* | *Reported by 507 (97%)* | *Reported by 345 (99%)* |
| American Indian or Alaskan Native | 6 (1.2%) | 6 (1.7%) |
| Black or African American | 23 (4.5%) | 21 (6.1%) |
| Hispanic or Latinx | 53 (11%) | 55 (16%) |
| East Asian | 102 (20%) | 65 (19%) |
| South Asian | 96 (19%) | 54 (16%) |
| Middle Eastern or North African | 16 (3.2%) | 14 (4.1%) |
| Native Hawaiian or Pacific Islander | 8 (1.6%) | 3 (0.9%) |
| White | 254 (50%) | 192 (56%) |
| *Gender* | *Reported by 516 (99%)* | *Reported by 343 (99%)* |
| Man | 149 (28.9%) | 89 (26%) |
| Woman | 362 (70.2%) | 254 (74%) |
| Another gender identity | 3 non-binary; 2 gender fluid (1.0%) | 1 non-binary; 1 gender non-conforming (0.6%) |
| *Parental Education* | *516 (99%)* | *343 (99%)* |
| No parents with a 4-yr degree | 116 (22.5%) | 75 (22%) |
| At least one parent with a 4-yr degree | 146 (28.3%) | 91 (27%) |
| At least one parent with a graduate degree | 254 (49.2%) | 182 (53%) |
| *Discipline* | *Reported by 511 (98%)* | *Reported by 345 (99%)* |
| Life Sciences | 356 (69.7%) | 245 (71%) |
| Chemistry | 53 (10.4%) | 35 (10%) |
| Engineering / Computer Science | 51 (10.0%) | 23 (6.7%) |
| Physics | 9 (1.8%) | 11 (3.1%) |
| Geosciences | 7 (1.4%) | 8 (2.3%) |
| Social Sciences | 16 (3.1%) | 14 (4.1%) |
| Allied Health | 8 (1.6%) | 3 (0.9%) |
| Interdisciplinary STEM | 9 (1.8%) | 4 (1.2%) |
| Math | 2 (0.4%) | 2 (0.6%) |
| *Mentor's Position* | *Reported by 518 (99%)* | *Reported by 343 (99%)* |
| Faculty | 325 (62.7%) | 197 (57%) |
| Postdoctoral associate | 68 (13.1%) | 38 (11%) |
| Graduate student | 84 (16.2%) | 84 (24%) |
| Undergraduate student | 26 (5.0%) | 24 (7.0%) |
| *Prior Research Experience* | *Reported by 515 (99%)* | NA |
| None | 18 (3.5%) |  |
| 1 term | 108 (21.0%) |  |
| 2 terms | 122 (23.7%) |  |
| 3 terms | 66 (12.8%) |  |
| More than 3 terms | 201 (39.0%) |  |

#### Factor Analysis

To examine the dimensionality of the MURS, we estimated an eight-factor confirmatory factor model for ordinal indicators using diagonally weighted least squares estimation in the ‘lavaan’ package (Rosseel, 2012) in R (R Core Team, 2021). Confirmatory Factor Analysis (CFA) is appropriate for established measures or when there is a theoretically-grounded reason to hypothesize about the factor structure. In contrast, Exploratory Factor Analysis is appropriate when the researchers do not have *a priori* hypotheses about the factor structure of the items. Because we had theorized dimensions during our qualitative study and the substantive phase, we had *a priori* expectations about the factor structure. Therefore, CFA is a more useful analytic strategy because it allowed us to test our hypothesized factor structure. We evaluated model fit holistically by considering both absolute and incremental indicators of fit: Root Mean Square Error of Approximation (RMSEA), Square Root Mean Residual (SRMR), Tucker-Lewis Index (TLI), and Comparative Fit Index (CFI).

Given that *inaccessibility*, *insufficient career and technical support*, *unclear and unreasonable expectations,* and *insufficient psychosocial support* were all measured by positively-phrased items, we reverse-scored them for analysis. We avoided having items with opposite valences within the same dimension to avoid introducing construct-irrelevant variance (e.g., error due to respondents misreading a negatively-worded item) (Roszkowski & Soven, 2010). We also recoded all items belonging to the *sexual harassment* factor to be dichotomous (0=Never; 1=Any frequency greater than “Never”) due to very low endorsement rates (95-98% of respondents chose “Never” for these items).

This model showed good fit to the data, c^2^(1511)=2,868.95, *p*<0.001, RMSEA=0.044 (95% CI: 0.042; 0.047), CFI=0.97, TLI=0.97, SRMSR=0.075, but resulted in two non-admissible solution problems. First, the correlation between *career & technical support* and *clear & reasonable expectations* factors was estimated at 0.96, causing the latent variable covariance matrix to be not positive definite (i.e., at least one factor in the model could be fully explained by a linear combination of the other factors). Second, the loading for one *sexual harassment* item (“My mentor made sexual comments about me”) was estimated at 1.02, resulting in a negative error variance. To resolve these problems, we collapsed the *career & technical support* and *clear & reasonable expectations* factors into a single factor, and deleted the item with a loading greater than 1.0. The resulting 7-factor model showed similar model-data fit, c^2^(1463)=2,922.69, *p*<0.001, RMSEA=0.047 (95% CI: 0.044; 0.0498), CFI=0.97, TLI=0.96, SRMSR=0.074. One item from the *abusive treatment* factor was found to have a loading less than 0.40 (l=0.36), and this item was deleted. After deletion, fit was similar, c^2^(1409)=2810.51, *p*<0.001, RMSEA=0.046 (95% CI: 0.044; 0.049), CFI=0.97, TLI=0.97, SRMSR=0.068.

The addition of the expected higher-order factors, supportive and destructive, worsened model-data fit, c^2^(1422)=3402.56, *p*<.001, RMSEA=.055 (95% CI: .053; .057), CFI=.95, TLI=.95, SRMSR=.083. To further inspect this issue, we analyzed the covariance matrix of the second-order latent variables using exploratory methods. The scree plot (see Supplemental Materials) suggested either two or three factors, as did other indicators of factor structure. Specifically, the Very Simple Structure (VSS) statistics, Velicer’s Minimum Average Partial (MAP) test, and Empirical Bayesian Information Criterion (BIC) suggested two factors whereas the sample-size-adjusted BIC suggested three^1^. Therefore, we examined both solutions. The two-factor model suggested the hypothesized two-factor structure was largely supported with one exception: the *interpersonal mismatch* factor cross-loaded onto both higher-order factors. The three-factor model suggested that *interpersonal mismatch* was a higher-order factor unto itself. To get more precise estimates we respecified our higher-confirmatory factor model for ordinal responses to include the cross-loading (Model A), and another model which specified *interpersonal mismatch* as its own factor (Model B). Model A showed good fit to the data, c^2^(1421)=3039.87, *p*<0.001, RMSEA=0.050 (95% CI: 0.047; 0.052), CFI=0.96, TLI=0.96, SRMSR=0.078, suggesting similar fit as the 7-factor model, but with a simpler model. The *interpersonal mismatch* factor loaded 0.58 on the **destructive** factor, and 0.44 onto the **supportive** factor. Model B yielded identical fit and *df* as in Model A but suggested a 0.91 correlation between the **destructive** and *interpersonal mismatch* factors, and therefore we proceed with the model including cross-loadings for the two higher-order factors as our final model. The higher-order structure of Model A, for which loadings were generally high, is shown in Figure 1. Similarly, first-order loadings ranged from 0.52 to 0.99, with a mean loading of 0.85, *SD=*0.11 (Table 2).

**Figure 1.**
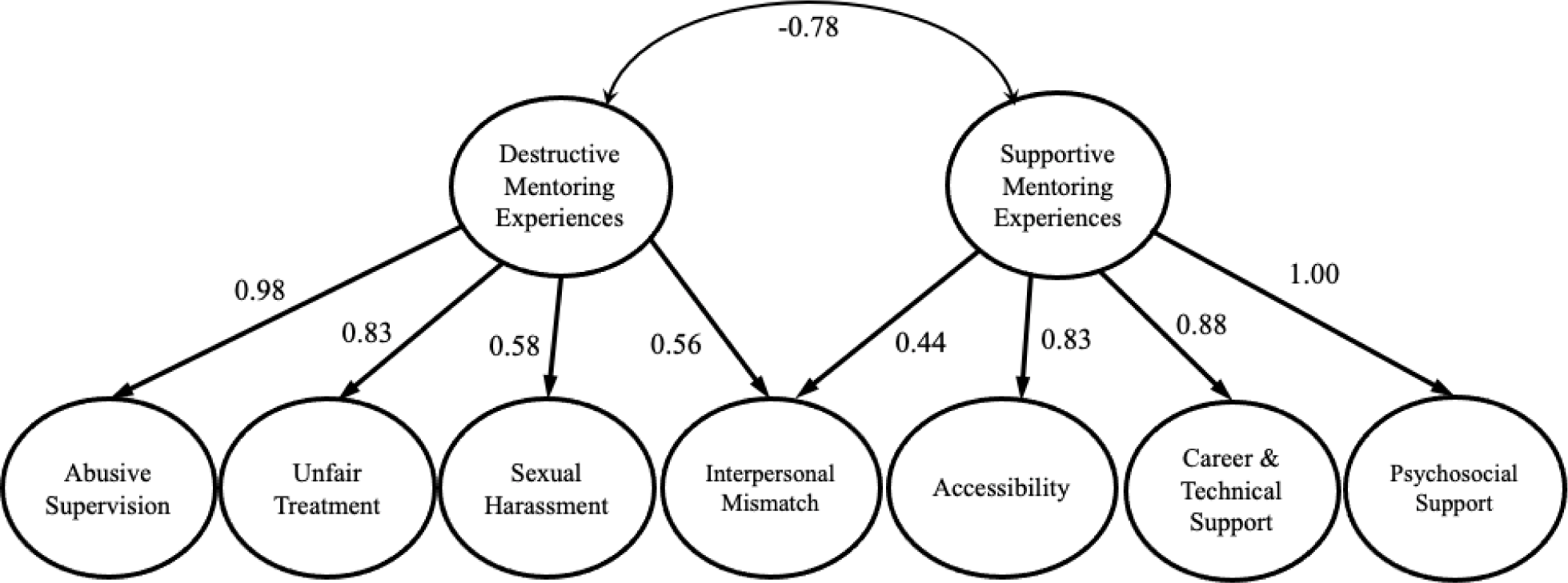
Final factor model for the Mentoring in Undergraduate Research Scale (MURS). Ovals represent latent factors and straight lines represent factor loadings. The final model includes seven first- order factors and two second-order factors. The first-order (item level) loadings are not pictured due to space, and can be found in Table 3. The negative correlation between the second order factors reflects the positive valence of supportive mentoring experiences and the negative valence of destructive mentoring experiences.

**Table 2.** Standardized first-order factor loadings for final MURS item set.

| **Dimension** | **Item** | **Standardized Loading ($\lambda$)** |
| --- | --- | --- |
| **Abusive supervision** |  |  |
| 1 | My mentor was rude to me. | 0.908 |
| 2 | My mentor belittled me. | 0.885 |
| 3 | My mentor created an intimidating environment. | 0.873 |
| 4 | My mentor was too harsh with their criticism. | 0.88 |
| 5 | My mentor was passive aggressive. | 0.879 |
| 6 | My mentor made me do excessive grunt work. | 0.712 |
| 7 | My mentor made inappropriate comments about my personal life. | 0.852 |
| 8 | My mentor was condescending. | 0.923 |
| 9 | My mentor blamed me for their mistakes. | 0.860 |
| **Accessibility (reverse-scored)** |  |  |
| 1 | My mentor gave me the attention I needed. | 0.946 |
| 2 | My mentor was available when I needed them. | 0.884 |
| 3 | My mentor was around to answer questions. | 0.870 |
| 4 | My mentor made time to meet with me. | 0.817 |
| 5 | My mentor responded when I contacted them. | 0.715 |
| **Career & Technical Support (reverse-scored)** |  |  |
| 1 | My mentor helped me understand the purpose of research tasks. | 0.763 |
| 2 | My mentor gave me work that was the right level of difficulty for me. | 0.702 |
| 3 | My mentor was clear about how my performance was being evaluated. | 0.657 |
| 4 | My mentor gave me the right amount of work. | 0.763 |
| 5 | My mentor made sure I was prepared to do research tasks. | 0.824 |
| 6 | My mentor gave me enough guidance in my research. | 0.840 |
| 7 | My mentor gave me useful feedback on my work. | 0.837 |
| 8 | My mentor was clear about what they wanted me to do. | 0.806 |
| **Psychosocial Support (reverse-scored)** |  |  |
| 1 | My mentor was friendly. | 0.87 |
| 2 | My mentor respected me. | 0.913 |
| 3 | My mentor had faith in me. | 0.842 |
| 4 | My mentor valued my contributions to the research. | 0.832 |
| 5 | My mentor cared about me as a person. | 0.847 |
| 6 | My mentor encouraged me. | 0.847 |
| **Interpersonal Mismatch** |  |  |
| 1 | My mentor and I had a tense relationship. | 0.899 |
| 2 | My mentor and I had incompatible personalities. | 0.872 |
| 3 | My mentor and I worked poorly together. | 0.92 |
| 4 | My mentor and I had difficulty getting along. | 0.934 |
| 5 | My mentor and I had incompatible communication styles. | 0.876 |
| **Sexual Harassment** |  |  |
| 1 | My mentor touched me without my permission. | 0.980 |
| 2 | My mentor made sexual remarks. | 0.917 |
| 3 | My mentor made sexual jokes. | 0.987 |
| **Unfair Treatment** |  |  |
| 1 | My mentor treated people unfairly based on their race/ethnicity | 0.89 |
| 2 | My mentor treated people unfairly based on their gender/sex | 0.892 |
| 3 | My mentor treated people unfairly based on their religion | 0.934 |
| 4 | My mentor treated people unfairly based on their sexual orientation | 0.958 |

**Table 3.** Item Response Theory (IRT) Model-Data Fit and Reliability Statistics. Model-data fit for each dimension has to be done one dimension at a time for IRT. M_2_ is the theoretical equivalent of Chi Square for IRT global fit; it is a Chi Square type statistic for nominal data, as opposed to continuous data. Rho xx’ is IRT-based marginal reliability. Misfit rate is the number of items showing significant misfit out of the total number of items retained from the CFA.

| Dimension | $M_2$ | $df$ | $p$ | RMSEA | 90% CI | | TLI | CFI | SRMR | $\rho_{xx'}$ | Misfit Rate |
| --- | --- | --- | --- | --- | --- | --- | --- | --- | --- | --- | --- |
|  |  |  |  |  | Low | High |  |  |  |  |  |
| Abusive Supervision | 96.33 | 35 | <0.001 | 0.059 | 0.045 | .074 | .95 | .96 | .066 | .86 | 0/14 |
| Inaccessibility | 32.05 | 5 | <0.001 | 0.102 | 0.070 | .137 | .97 | .98 | .042 | .87 | 0/5 |
| Insufficient Career & Technical Support | 232.69 | 54 | <0.001 | 0.080 | 0.069 | .091 | .96 | .97 | .054 | .91 | 0/12 |
| Insufficient Psychosocial Support | 229.47 | 14 | <0.001 | 0.174 | 0.154 | .194 | .08 | .92 | .940 | .87 | 0/7 |
| Interpersonal Mismatch | 68.26 | 9 | <0.001 | 0.113 | 0.088 | .139 | .97 | .98 | .064 | .85 | 0/6 |
| Sexual Harassment | 5.73 | 2 | 0.057 | 0.060 | 0.000 | .121 | .98 | .99 | .083 | .16* | n/a |
| Unfair Treatment | 120.72 | 14 | <0.001 | 0.123 | 0.103 | .144 | .96 | .97 | .059 | .72 | 1/8<sup>+</sup> |

RMSEA= Root Mean Square Error of Approximation; TLI = Tucker-Lewis Index; CFI=Comparative Fit Index; SRMR=Standardized Root Mean Residual; $r_{xx'}$ =IRT-based marginal reliability; Misfit Rate = The number of items with significant misfit according to the $s-\chi^2$ item fit statistic out of the total number of items. <sup>+</sup> Indicates the misfitting item was removed prior to estimating global model-data fit. <sup>\*</sup> Although the overall marginal reliability of the sexual harassment scale was low, its reliability at high levels of the construct was acceptable. Global fit and marginal reliability were estimated prior to items being deleted unless noted otherwise.

We conducted item response theory analyses to further refine the MURS subscales by removing items that did not contribute to the reliability of the measure. For each subscale, we estimated the graded response model (GRM; (Samejima, 1968)). Global (i.e., scale-level) model-data fit was evaluated using the family of *M*_2_-based goodness of fit statistics (see Cai & Hansen, 2013; Cai & Monroe, 2014; Maydeu-Olivares & Joe, 2006). The *M*_2_ statistic is statistically equivalent to the c2 used in structural equation modeling; its properties allow for calculating fit indices such as the RMSEA, CFI, and TLI. At the item-level, model-data fit was assessed using the *S-*χ^2^ statistic (Orlando & Thissen, 2000), which examines the degree to which observed item responses deviate from expectations across the distribution of the latent variable. All analyses were conducted in the ‘mirt’ package (Chalmers, 2012). Table 3 shows the results of the model-data fit analyses and IRT-based marginal reliability for each scale. In general, these results suggested good model-data fit, which indicates item parameters and the resulting information functions are stable and interpretable.

Given the good model-data fit, we continued to examine each subscale and remove items that provided very low information. For *abusive supervision*, we found that five of 14 items had very low information across the construct continuum (My mentor gossiped about people in the lab; scolded people in the lab; invaded my privacy; discussed topics that were too personal; and took credit for my work). The *inaccessibility* measure showed none of its five items that required removal. The *career & technical support* scale showed four of twelve items that had very low information (My mentor explained how my work fit into the bigger picture; was clear about when I was expected to be working; expected me to work reasonable hours; and my mentor and I talked about my career aspirations). Only one item in the *psychosocial support* scale showed low information (My mentor thought the work I did was important). For the *interpersonal mismatch* scale, only one item (My mentor and I had incompatible work styles) showed low information. Of the 8 items for *unfair treatment*, one was found to have extremely high misfit (My mentor treated people unfairly based on their career interests) and was removed prior to estimating global model-data fit. The remaining 7 item measure showed low information for three items (My mentor treated people unfairly based on their major; was biased against certain groups of people, and had favorites in the lab).

Given that only three items were included in the *sexual harassment* measure, we added “My mentor made sexual comments about me” back to the item set to achieve model identification. The fit was very good, but the marginal reliability was very low at 0.16; this is due to the fact that it measures such extreme and rare behavior (e.g., touching without permission, sexual remarks) that it only has high reliability. In other words, at 2 *SD* above the mean the IRT reliability of the scale is 0.96, but the measure is low in reliability for those near the mean. The second and third items are repetitive and are extremely highly correlated, and thus only one could be used (the item “My mentor made sexual comments about me” caused a loading greater than 1.0 in our initial model for the same reason), as their content is highly similar (i.e., making sexual remarks versus making sexual remarks about the respondent specifically). The final MURS items and their standardized factor loadings are presented in Table 2.

#### Personality and negative mentoring

We examined correlations between the Big Five personality traits (openness, conscientiousness, extraversion, agreeableness, neuroticism) and the MURS factors. Our aim was to ensure that the MURS was not measuring facets of personality that might make individuals more or less likely to report negative experiences with mentors. Openness was the only personality trait that significantly related to negative mentoring, showing a weak negative association (*r* = -0.14, *p* = 0.001). Openness also exhibited significant but small negative associations with most dimensions of negative mentoring: *abusive supervision* (*r* = -0.12, *p* = 0.006), *inaccessibility* (*r* = -0.12, *p* = 0.008), *insufficient career and technical support* (*r* = -0.13, *p* = 0.004), *insufficient psychosocial support* (*r* = -0.10, *p* = 0.02), *interpersonal mismatch* (*r* = -0.14, *p* = 0.001), *sexual harassment* (*r* = -0.08, *p* = 0.06), *unfair treatment* (*r* = -0.06, *p* = 0.19). These results suggested that students’ perceptions of their mentoring experiences may be broadly influenced by their level of openness. To account for this, we opted to measure openness in the next phase of data collection so that we could ensure that it did not influence the outcomes of interest and confound our ability to estimate the impact of negative mentoring experiences. We also chose to measure neuroticism to ensure the replicability of the lack of association with negative mentoring experiences.

### External Phase

In our final phase of data collection and analysis, we aimed to characterize relationships between responses on the MURS with variables we hypothesized would relate to negative mentoring experiences, or its nomological network (Cronbach & Meehl, 1955). In other words, we sought to interpret the meaning of the MURS scores in relation to theoretically-relevant constructs and outcomes (Benson, 1998). To accomplish this, we collected evidence of the discriminant, convergent, and predictive validity of the MURS. Scores on the MURS should be correlated with measures of related constructs (i.e., convergent validity), should not correlate with measures of unrelated constructs (i.e., discriminant validity), and should be predictive of theoretically-related and practically-relevant outcomes. For ease of reading, we present the methods for data collection and analysis first. Then, we describe our hypotheses of how responses on the MURS relate to student outcomes, covariates, and other measures of mentoring quality along with our results characterizing these relationships. We continue to present methods and results using a common, negative valence for ease of reading and comparison.

#### Data Collection

To carry out the external phase, we collected and analyzed a second national dataset. We recruited undergraduates at 32 institutions who were about to do research for the *first time* to avoid selection bias in the sample (i.e., students staying or leaving research experiences because of the mentorship they experienced). We did not include any selective programs (i.e., programs that had an application process or selected students based on academic standing, such as honors programs and REU programs) to mitigate bias in our sample. We used a pre/post-survey design to evaluate how students’ negative mentoring experiences related to changes they may or may not realize from participating in undergraduate research.

Prior to the start of their research experience, we emailed participants a pre-survey with measures of our constructs of interest as well as items to measure student demographics (see full item set with references and description of validity evidence in Supplemental Material). At the end of one term of research (quarter, semester, summer), we emailed them the post-survey, which included the MURS items along with measures of our constructs of interest (see full item set with references and description of validity evidence in Supplemental Material). Students were compensated $25 total for their participation: $10 for the pre-survey, $15 for the post-survey. We received 359 responses to both the pre- and post-survey; 11 of which did not pass all attention checks. Thus, the final sample size was n=348 (Table 1).

#### Base Rate of Mentoring Experiences

We plotted histograms (Figure 2) and calculated means and standard errors (SE; Table 4) to gain insight into prevalence of mentoring experiences. Undergraduates reported the highest absence of *career and technical support* (M = 1.55, SD = 0.72) compared to all other forms of negative mentoring, although its base rate was still low (response scale was 1 to 5, with higher values indicating more negative mentoring experiences). Undergraduates reported lower levels of *abusive supervision* (M = 1.23, SD = 0.49), *inaccessibility* (M = 1.32, SD = 0.61), *insufficient psychosocial support* (M = 1.30, SD = 0.45), *interpersonal mismatch* (M = 1.27, SD = 0.57) and the lowest levels of *sexual harassment* (m = 1.02, SD = 0.10) and *unfair treatment* (M = 1.09, SD = 0.51), indicating that these forms of negative mentoring were quite uncommon in our sample.

**Figure 2.**
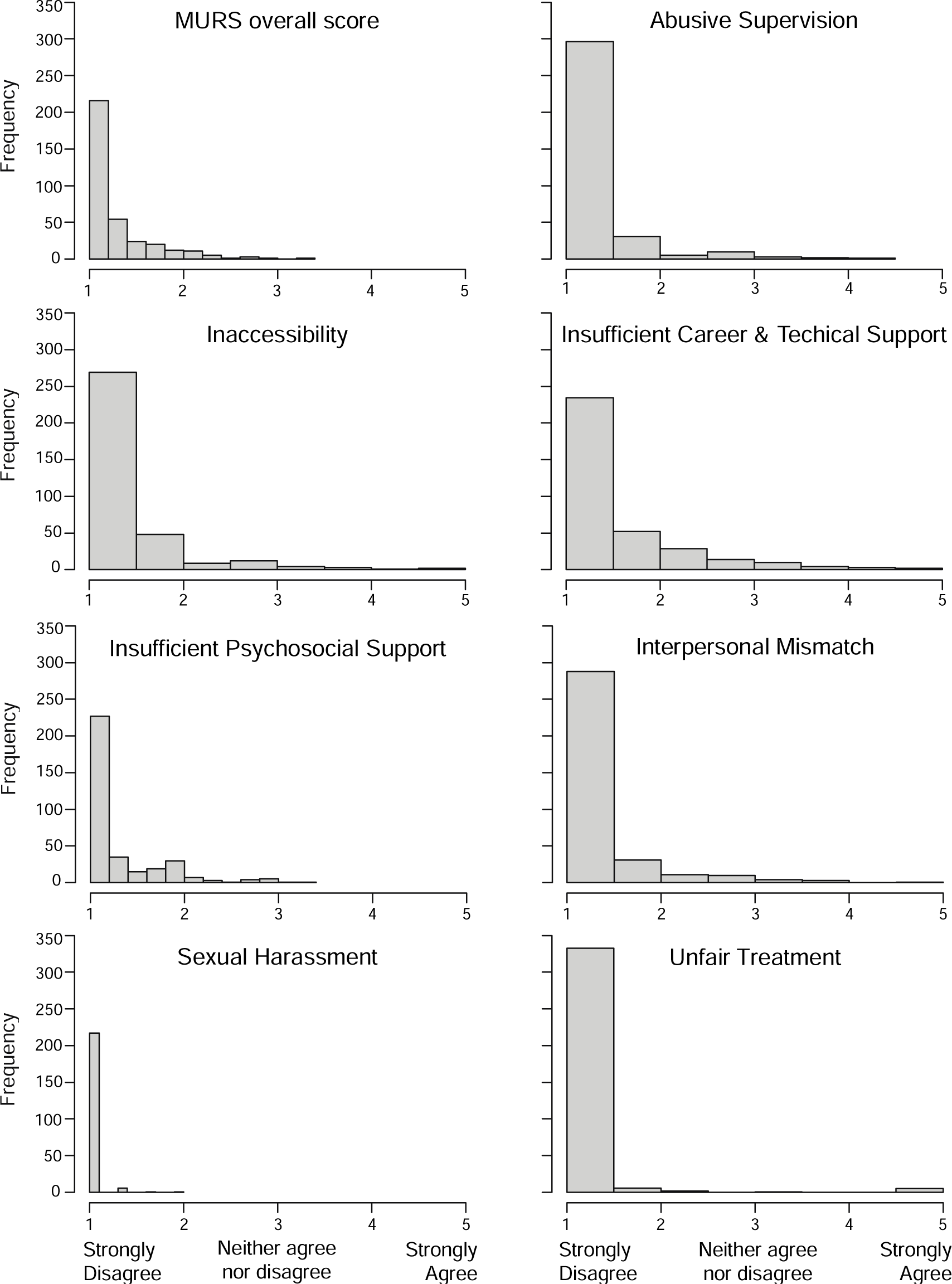
Histograms of the MURS and its seven dimensions. For comprehensibility and ease of comparison, histograms reflect our use of a common, negative valence such that supportive experience results are reverse scored and presented in terms of their absence (inaccessibility, insufficient career & technical support, insufficient psychosocial support) and destructive experiences are presented in terms of their presence. The overall MURS score reflects negative mentoring experiences.

**Table 4.** Descriptive statistics and correlations for external phase data. Relationships that were predicted a priori are indicated with the following emphases: shading indicate a predicted negative relationship and a double border ⎜⎜indicates a predicted positive relationship. Significant relationships are **bolded** with \**p* < 0.05, \*\**p* < 0.01, \*\*\**p* < 0.001.

| Construct & Variable |  | N | M | SD | Abusive Supervision | Inaccessibility | Insufficient Career & Technical Support | Insufficient Psychosocial Support | Interpersonal Mismatch | Sexual Harassment | Unfair Treatment | MURS overall |
| --- | --- | --- | --- | --- | --- | --- | --- | --- | --- | --- | --- | --- |
| Negative Mentoring | Abusive Supervision | 348 | 1.23 | 0.49 |  |  |  |  |  |  |  |  |
|  | Inaccessibility | 348 | 1.32 | 0.61 | **0.44\*\*\*** |  |  |  |  |  |  |  |
|  | Insufficient Career & Technical Support | 348 | 1.55 | 0.72 | **0.39\*\*\*** | **0.70\*\*\*** |  |  |  |  |  |  |
|  | Insufficient Psychosocial Support | 348 | 1.30 | 0.45 | **0.58\*\*\*** | **0.60\*\*\*** | **0.67\*\*\*** |  |  |  |  |  |
|  | Interpersonal Mismatch | 348 | 1.27 | 0.57 | **0.72\*\*\*** | **0.53\*\*\*** | **0.54\*\*\*** | **0.64\*\*\*** |  |  |  |  |
|  | Sexual Harassment | 225 | 1.02 | 0.10 | 0.08 | -0.01 | **0.14\*** | 0.06 | 0.03 |  |  |  |
|  | Unfair Treatment | 347 | 1.09 | 0.51 | 0.07 | 0.03 | 0.03 | 0.04 | 0.10 | 0.08 |  |  |
| Personality | Neuroticism | 348 | 3.34 | 0.96 | -0.01 | -0.05 | -0.04 | -0.03 | 0.01 | -0.03 | -0.01 | -0.03 |
|  | Openness | 348 | 3.85 | 0.87 | -0.04 | -0.02 | 0.00 | -0.06 | -0.01 | 0.13 | 0.00 | -0.03 |
| Attachment Style | Avoidant | 348 | 2.52 | 0.89 | 0.02 | 0.06 | 0.10 | 0.09 | 0.09 | 0.07 | 0.06 | 0.10 |
|  | Anxious | 348 | 2.72 | 0.90 | 0.06 | **0.16\*\*** | **0.17\*\*** | **0.12\*** | **0.16\*\*** | **0.16\*** | -0.06 | **0.15\*\*** |
|  | Secure | 348 | 4.28 | 0.68 | -0.09 | -0.06 | -0.08 | **-0.12\*** | -0.08 | 0.11 | 0.00 | -0.10 |
| Emotions | Positive Affect Toward Research | 341 | 4.02 | 0.88 | **-0.17\*\*** | **-0.31\*\*\*** | **-0.46\*\*\*** | **-0.42\*\*\*** | **-0.28\*\*\*** | -0.12 | -0.02 | **-0.39\*\*** |
|  | Negative Affect Toward Research | 341 | 2.12 | 0.70 | **0.43\*\*\*** | **0.24\*\*\*** | **0.33\*\*\*** | **0.30\*\*\*** | **0.45\*\*\*** | 0.05 | 0.08 | **0.42\*\*\*** |

**Table 4. Descriptive statistics and correlations for external phase data.**
| Construct & Variable |  | N | M | SD | Abusive Supervision | Inaccessibility | Insufficient Career & Technical Support | Insufficient Psychosocial Support | Interpersonal Mismatch | Sexual Harassment | Unfair Treatment | MURS overall | Mentoring Competence |
| --- | --- | --- | --- | --- | --- | --- | --- | --- | --- | --- | --- | --- | --- |
| Mentoring Competence |  | 347 | 3.64 | 0.48 | **-0.46\*\*\*** | **-0.62\*\*\*** | **-0.79\*\*\*** | **-0.70\*\*\*** | **-0.56\*\*\*** | -0.10 | -0.08 | **-0.75\*\*\*** |  |
| Mentoring Relationship Quality |  | 340 | 4.47 | 0.80 | **-0.45\*\*\*** | **-0.73\*\*\*** | **-0.77\*\*\*** | **-0.72\*\*\*** | **-0.59\*\*\*** | -0.06 | 0.00 | **-0.76\*\*\*** | **0.78\*\*\*** |
| Scientific self-efficacy | Pre | 348 | 3.50 | 0.68 | 0.05 | -0.08 | **-0.11\*** | -0.10 | -0.07 | 0.01 | **-0.14\*** | -0.10 |  |
|  | Post | 340 | 4.12 | 0.64 | **-0.13\*** | **-0.31\*\*\*** | **-0.39\*\*** | **-0.37\*\*\*** | **-0.23\*\*\*** | -0.02 | -0.06 | **-0.34\*\*\*** |  |
| Scientific identity | Pre | 348 | 4.06 | 0.67 | -0.03 | **-0.16\*\*** | **-0.23\*\*\*** | **-0.21\*\*\*** | **-0.12\*** | 0.01 | -0.05 | **-0.19\*\*\*** |  |
|  | Post | 340 | 4.24 | 0.76 | **-0.12\*** | **-0.27\*\*** | **-0.38\*\*\*** | **-0.38\*\*\*** | **-0.21\*\*\*** | -0.05 | 0 | **-0.32\*\*\*** |  |
| Research beliefs | Pre | 348 | 3.98 | 0.48 | **-0.18\*\*\*** | -0.08 | **-0.22\*\*\*** | **-0.18\*\*\*** | **-0.23\*\*\*** | -0.04 | -0.04 | **-0.23\*\*\*** |  |
|  | Post | 341 | 3.98 | 0.59 | **-0.25\*\*\*** | **-0.22\*\*\*** | **-0.33\*\*\*** | **-0.27\*\*\*** | **-0.29\*\*\*** | -0.07 | -0.09 | **-0.34\*\*\*** |  |
| Intentions | Pre | 347 | 4.19 | 0.68 | **-0.15\*\*** | -0.08 | **-0.18\*\*\*** | **-0.16\*\*** | **-0.14\*\*** | -0.09 | 0.03 | **-0.16\*\*** |  |
|  | Post | 341 | 4.15 | 0.84 | **-0.15\*\*** | **-0.19\*\*\*** | **-0.23\*\*\*** | **-0.17\*\*** | **-0.16\*\*** | -0.09 | -0.03 | **-0.22\*\*\*** |  |

We looked for differences in students’ reports of mentoring experiences based on their personal characteristics using Wilcoxon Rank-Sum tests (i.e., non-parametric t-tests) and Kruskal-Wallis tests (i.e., non-parametric ANOVAs). We found no differences in any dimensions of mentoring by race/ethnicity when comparing the experiences of students who identified as Asian, White, or from a minoritized race or ethnicity^2^. We also found no differences by generation in college, or by mentor rank (faculty or not faculty). We only observed one difference by student gender: men reported more *abusive supervision* than women (men: n=89, M = 1.35, SD 0.63; women: n=254, M = 1.19, SD = 0.43, W = 13392, *p* = 0.003).

#### Discriminant and Convergent Validity

We evaluated discriminant and convergent validity by making *a priori* predictions about how mentoring experiences would relate to students’ personality traits, attachment styles, emotions about research, and other measures of mentoring. We then evaluated these relations by examining bivariate correlations (Table 4). Given the exploratory nature of this work, we opted to use a less conservative cut-off value of *p* < 0.05 to determine significance and we report all *p* values for readers to make their own judgments.

#### Personality traits

We hypothesized that mentoring experiences would be unrelated to any personality traits. Based on results from the structural phase analysis, we sought to rule out the hypothesis that an undergraduate’s reports of negative mentoring experiences were due to their level of openness. Prior research indicates that individuals high on openness are likely to judge experiences as less negative and more likely to respond to abusive supervision with coping strategies that mitigate the emotional labor associated with such experiences (Steel et al., 2008; Wu & Hu, 2013). Given that the personality trait of neuroticism reflects a tendency toward negative feelings, we also sought to rule out the hypothesis that an undergraduate’s reports of negative mentoring experiences were due to their level of neuroticism. As predicted, all correlations between the seven negative mentoring dimensions and both *neuroticism* and *openness* were near-zero and non-significant (Table 4). These results suggest that the MURS is unlikely to be measuring personality traits per se and provide evidence of discriminant validity.

#### Attachment styles

Attachment styles are stable patterns of emotions and behaviors exhibited in close relationships, which are thought to develop through early interactions between infants and their caregivers (Ainsworth, 1989; Bowlby, 1979; Bowlby & Ainsworth, 2013). Researchers have described two main forms of attachment: *secure*, in which the infant perceives their caregiver as a source of comfort and strength, and insecure or anxious attachment. Forms of insecure attachment include *anxious* attachment, in which the infant perceives their caregiver as an unreliable – sometimes offering support and other times not, and *avoidant* attachment, in which the infant has learned the caregiver is not a reliable source of support and thus does not expect or seek comfort from them (Carver, 1997). These early experiences are thought to shape an individual’s internal working model of relationships and thus influence how adults think, feel, and behave in close relationships (Hazan & Shaver, 1994), including supervisory relationships (Fitch et al., 2010). Thus, we sought to explore whether and how undergraduate researchers’ attachment styles related to their negative mentoring experiences.

We hypothesized that undergraduates’ mentoring experiences would relate to their attachment styles, but that the magnitude of the correlations would be small to moderate such that mentoring experiences are not redundant with attachment style. We focused on avoidant and anxious attachment styles because we hypothesized that these attachment styles would influence an undergraduate researcher’s expectations for their relationships with their mentors. Specifically, we predicted that *avoidant* attachment style would negatively relate to both *inaccessibility* and *insufficient psychosocial support* because individuals who are avoidant would expect less attention and support from their mentors having learned to not expect such support from their caregivers. Thus, they would be less likely to report dissatisfaction when their mentor was unavailable to them or did not provide psychosocial support. However, we found that undergraduates’ levels of *avoidant* attachment were not associated with their ratings of mentor *inaccessibility* (*r* = 0.06, *p* = 0.29) or *insufficient psychosocial support* (*r* = 0.09, *p* = 0.093) (Table 4).

Research indicates that anxious attachment includes an individual’s fear of abandonment in relationships as well as the tendency to want to have a closer relationship than their relational partner (Carver, 1997). We hypothesized that undergraduates’ levels of anxious attachment would positively relate to *inaccessibility* and *insufficient psychosocial support* because individuals with anxious attachment styles desire a higher level of attention and support and thus may be more distressed by these forms of negative mentoring. Surprisingly, undergraduates who indicated an *anxious* attachment style reported slightly higher levels of most forms of negative mentoring, except *abusive supervision* and *unfair treatment* (Table 4). We did not have *a priori* hypotheses about secure attachment style and negative mentoring experiences. Yet, we observed a small but significant relationship between undergraduates reporting a secure attachment style and lower levels of *insufficient psychosocial support* (*r* = -0.12, *p* < 0.05) (Table 4). Altogether, these results indicate that undergraduate researchers who have an *anxious* attachment style may be slightly more susceptible to negative mentoring experiences. Furthermore, undergraduates with a secure attachment style might perceive more psychosocial support or require less psychosocial support to thrive. Collectively, however, these effects were modest (*r* values from |0.10| to |0.17|; Table 4), which indicates that the MURS is unlikely to simply be measuring attachment styles, providing further evidence of discriminant validity.

#### Emotions about research

Emotions are responses, including feelings, actions, and physiological changes, to situations that garner an individual’s attention (Gross & Thompson, 2007). Appraisal theory indicates that emotions arise when an individual positively or negatively appraises a situation that is personally significant to them (Scherer, 1999). Prior research shows that students’ emotions can have substantial effects on their academic engagement and performance (Pekrun & Linnenbrink-Garcia, 2012). In addition, negative behaviors in the workplace are associated with employees experiencing toxic emotions and emotional exhaustion (e.g., Han et al., 2017; Henle & Gross, 2014; Porath & Pearson, 2012). Thus, we hypothesized that negative mentoring experiences would impact whether students have positive or negative emotions about their research experience. Specifically, we hypothesized that students’ positive emotions about research (e.g., excitement, accomplishment) would be negatively related to experiencing *insufficient career and technical support* and *insufficient psychosocial support* because we postulated that students who experience these forms of support are more likely to feel positively about themselves and their work. Indeed, undergraduates who reported higher levels of *insufficient career and technical support* and *insufficient psychosocial support*, as well as all other forms of negative mentoring except *sexual harassment* and *unfair treatment*, reported significantly lower levels of *positive emotions* (*r* values from - 0.17 to -0.46; Table 4). We also hypothesized that students’ *negative emotions* about research (e.g., stress, apathy) would be positively related to all forms of negative mentoring because all of these experiences are likely to generate mentee distress. As expected, undergraduates’ *negative emotions* about research were significantly correlated with all forms of negative mentoring experiences except *sexual harassment* and *unfair treatment* (*r* values ranged from 0.24 to 0.45; Table 4).

#### Other measures of mentoring

If the MURS is measuring the range of mentoring undergraduates experience, responses on the MURS should relate to the perceived quality of their mentoring relationships (Allen & Eby, 2003). Responses on the MURS should also relate to measures of perceived mentoring competency, including a mentor’s abilities to communicate effectively with their mentee, align their expectations with those of their mentee, and foster their mentee’s independence (Fleming, House, Shewakramani, et al., 2013). Specifically, we predicted that:

- *Mentoring relationship quality* will negatively relate to the overall MURS score and to *all seven dimensions* of negative mentoring because negative mentoring experiences should undermine the development and maintenance of a quality relationship.
- *Mentoring competence* will negatively relate to the overall MURS score and to *all seven dimensions* of negative mentoring because mentoring competence is needed to prevent negative mentoring experiences.

Undergraduates who reported lower levels of mentoring relationship quality reported significantly higher levels of negative mentoring experience overall (*r* = -0.75, *p* < 0.001) and of all dimensions of negative mentoring except *sexual harassment* and *unfair treatment* (*r* values ranged from -0.46 to -0.79; Table 4). Undergraduates who reported lower levels of mentor competence also reported significantly higher levels of negative mentoring overall (*r* = -0.76, *p* < 0.001) and of all dimensions of negative mentoring, except *sexual harassment* and *unfair treatment* (*r* values from -0.45 to -0.77; Table 4). Collectively, these results indicate that the MURS is measuring aspects of mentoring relationships that relate to mentoring quality and mentor competency, without being completely redundant with these measures.

#### Predictive Validity

Finally, we examined how mentoring experiences measured by MURS related to outcomes undergraduates typically experience from participating in research. Research experiences are widely accepted as formative experiences in which undergraduates grow in their belief that they can be successful in science (i.e., science self-efficacy) and their view of themselves as a “science person” (i.e., scientific identity) (Estrada et al., 2011; Gentile et al., 2017; Hunter et al., 2007; Kardash, 2000; Robnett et al., 2015). Furthermore, expectancy value theory postulates that one is motivated to engage in a task, such as pursuing a science research career, if one believes they can be successful (i.e., science self-efficacy) and that the task has value (e.g., the benefits of doing science research outweigh the costs) (Barron & Hulleman, 2015; Wigfield & Eccles, 2000). Based on this research and theory, we formulated a series of hypotheses regarding how experiencing negative mentoring would relate to undergraduate researchers’ development of *science self-efficacy* and *science identity* as well as their beliefs about the value of research (*research beliefs*) and their intentions to continue in science and in research (*intentions*). We evaluated these relationships by fitting a series of linear regression models using mean-scores for relevant scales, as in this example model: Outcome_t2 ∼ Outcome_t1 + MURS We sought to determine whether MURS explained variance in undergraduates’ post-research self-efficacy, identity, beliefs, and intentions above and beyond their pre-research ratings (Table 5).

**Table 5.** Regression analysis results. β = standardized estimate. ** *p* < 0.01, *** *p* < 0.001.

| Model & Predictors |  | Scientific Self-Efficacy Post-score |  | Scientific Identity Post-score |  | Research beliefs Post-score |  | Intentions Post-score |  |
| --- | --- | --- | --- | --- | --- | --- | --- | --- | --- |
| | | $\beta$ | Variance explained | $\beta$ | Variance explained | $\beta$ | Variance explained | $\beta$ | Variance explained |
| Baseline | Pre-score | 0.39*** | $R^2 = 0.15$ | 0.58*** | $R^2 = 0.33$ | 0.52*** | $R^2 = 0.27$ | 0.67*** | $R^2 = 0.45$ |
| MURS | Pre-score | 0.36*** | $R^2 = 0.25$ | 0.54*** | $R^2 = 0.38$ | 0.47*** | $R^2 = 0.32$ | 0.65*** | $R^2 = 0.46$ |
|  | MURS | -0.31*** |  | -0.22*** |  | -0.23*** |  | -0.13** |  |
| MCA | Pre-score | 0.36*** | $R^2 = 0.30$ | 0.51*** | $R^2 = 0.43$ | 0.48*** | $R^2 = 0.33$ | 0.65*** | $R^2 = 0.47$ |
|  | MCA | 0.38*** |  | 0.31*** |  | 0.24*** |  | 0.14*** |  |
| MRQ | Pre-score | 0.36*** | $R^2 = 0.27$ | 0.51*** | $R^2 = 0.45$ | 0.49*** | $R^2 = 0.31$ | 0.65*** | $R^2 = 0.46$ |
|  | MRQ | 0.35*** |  | 0.34*** |  | 0.20*** |  | 0.13** |  |

#### Scientific self-efficacy

Research indicates that social persuasion, meaning encouraging feedback from influential individuals, such as mentors, functions as a source of self-efficacy (Usher & Pajares, 2008). Undergraduate researchers may experience less development of their *scientific self-efficacy* if they do not experience social persuasion because their mentors are inaccessible or are not providing psychosocial support. Mastery experiences, or tackling and ultimately succeeding at a challenging task, function as another critical source of self-efficacy development (Usher & Pajares, 2008). Undergraduate researchers may have fewer mastery experiences if they receive insufficient support to be successful, their tasks are not at the right level of challenge, or they perceive themselves to be unsuccessful. Thus, we hypothesized that students’ development of *scientific self-efficacy* during research would be limited by experiencing negative mentoring. As expected, undergraduates reported significantly lower post-research self-efficacy when they reported experiencing more negative mentoring, even after accounting for their pre-research self-efficacy (β = -0.31, *p* < 0.001; Table 5).

#### Scientific identity

Typically, undergraduate research experiences positively influence students’ *scientific identity*, making them feel like more of a “science person” (Estrada et al., 2011; Gentile et al., 2017; Robnett et al., 2015). Identity development or lack thereof, is influenced by recognition from members of the community, such as mentors (Carlone & Johnson, 2007; Hazari et al., 2010). Thus, we hypothesized that students’ development of a *scientific identity* during research would be limited by experiencing negative mentoring. Indeed, undergraduates’ post-research *scientific identity* was limited significantly by experiencing more negative mentoring (β = -0.22, *p* < 0.001; Table 5).

#### Research beliefs

We hypothesized that undergraduates who experienced more negative mentoring would perceive research as less beneficial and more costly (Ceyhan & Tillotson, 2020; Gaspard, Dicke, Flunger, Brisson, et al., 2015; Gaspard, Dicke, Flunger, Schreier, et al., 2015). We focused on measuring students’ beliefs about the *intrinsic value* of research (i.e., how interesting or enjoyable research is), the *communal value* of research (i.e., potential for research to benefit a broader community or society), and the *opportunity costs* of research (i.e., sacrifices students perceive they would have to make to engage in research) (Barron & Hulleman, 2015; Brown et al., 2015). We hypothesized that, collectively, students’ post-research beliefs would be hampered by negative mentoring experiences (increased perceptions of opportunity costs and decreased perceptions of intrinsic and communal value). As expected, undergraduates’ post-research *beliefs* were limited significantly by experiencing more negative mentoring (β = -0.23, *p* < 0.001; Table 5).

#### Graduate and career intentions

Undergraduates who participate in research clarify their career choices and, as a result, can change their *intentions* to pursue graduate education and careers in science (Estrada et al., 2011, 2018; Gentile et al., 2017). We hypothesized that students’ *intentions* would negatively relate to experiencing more negative mentoring because, if students do not receive sufficient support, perceive a mismatch with more experienced researchers, or are treated poorly or unfairly, they are more likely to opt out of science or research paths. Indeed, undergraduates’ post-research *intentions* were limited significantly – although to a lesser extent than other outcomes – by experiencing more negative mentoring (β = -0.13, *p* < 0.01; Table 5).

We next examined correlations between dimensions of the MURS and pre- and post-measures of each outcome. As expected, all of the dimensions of MURS were negatively related to students’ post-research ratings of their self-efficacy, identity, research beliefs, and intentions, except for sexual harassment and unfair treatment. Surprisingly, students’ pre-research ratings of their identity, beliefs, and intentions were also negatively related to responses on the MURS, although these relationships were more modest (*r* values from -0.12 to -0.23). It may be that students who identify less as a scientist, who hold more skeptical beliefs about the value of doing research, or who do not have strong intentions to continue in science or research have greater mentoring needs and thus report less favorable mentoring experiences. Alternatively, mentors may be consciously or unconsciously sensing that their mentees are less integrated into the scientific community and proffering less favorable mentoring.

In order to compare the explanatory values of the MURS versus measures of mentoring relationship quality and mentoring competence, we fit a similar series of linear regression models using the mentoring relationship quality scale (MRQ) or the mentoring competency assessment (MCA), as in this example: Outcome _t2 ∼ Outcome_t1 + MRQ. The standardized estimates for the MURS, MRQ, and MCA were quite similar for all of the outcomes we examined (Table 5). In addition, the variance in outcomes explained by mentoring was similar, regardless of whether negative mentoring (MURS), mentoring relationship quality (MRQ), or mentoring competence (MCA) was the focus of analysis. In other words, all three mentoring measures explained variance in post-research self-efficacy, identity, beliefs, and intentions beyond pre-research ratings, but the variance explained was similar.

### Limitations

Given the potential for measurement tools to shape future research and findings, we took several steps to protect against threats to validity of MURS as a measure of mentoring experiences. We collected data from a diversity of undergraduate researchers at a variety of types of institutions. We collected data from students who varied in their gender, racial, and other identities, which bolsters the potential applicability of our results to diverse student groups. Yet, we were not able to collect sufficient responses to allow for examination of the experiences of particular groups of students (e.g., individuals identifying as Native American, Black, non-binary). Further research should be done to examine whether students who identify with particular groups experience negative mentoring at different rates, or are differentially affected by it.

While our results from the structural phase show strong evidence in support of the theorized factor structure, our external phase sample was too small to test the structural model fit identified from the first sample. Furthermore, the external phase sample was too small leverage the strength of structural equation modeling to examine the influence of various forms of negative mentoring on undergraduate researchers’ outcomes while accounting for measurement error. Future research should either examine particular dimensions of interest (rather than the whole measure), or collect data from a sufficiently large sample to estimate the measurement models.

Our study lacks validity evidence related to the consequences of testing (American Educational Research Association et al., 2014), which should be addressed in future work. For instance, stakeholders in undergraduate science education are likely to be interested in protecting undergraduate researchers from negative mentoring. They may administer the MURS, identify “negative mentors” based on scores from undergraduate researchers in their research group, and prevent those mentors from working with undergraduate researchers in the future. In other words, the scores on the MURS may lead to judgments about who can or cannot mentor. This interpretation may not be valid because mentorship is inherently dyadic and embedded in a context, such as a program or degree plan that exerts additional influence on the mentoring relationship. The mentees themselves and these other contextual factors could be contributing to dysfunction in mentoring relationships. Furthermore, mentees differ in their goals, interests, experiences, and aspirations and thus require different investments of time, training, and support from mentors. Thus, a mentor who may be a poor fit with one mentee may be an excellent fit with another. Finally, mentees themselves may be biased against particular mentors based on their identities, as has been observed in student end-of-course evaluations of instruction (Esarey & Valdes, 2020; Fan et al., 2019; Goos & Salomons, 2017; MacNell et al., 2015). Future research needs to examine and study potential unintended negative consequences of the MURS for mentees, mentors, and programs.

Our study had several limitations beyond those associated with validity. First, we conducted a large number of tests, which may have resulted in some false positives. Our intention was to explore and characterize relationships that offer insight into the validity and utility of the MURS. However, the relationships reported here should continue to be investigated in future research. Second, our results suggest that the sexual harassment and unfair treatment dimensions of the MURS need additional refinement. We failed to find the relationships between these forms of negative mentoring and almost all of the other constructs we hypothesized to be related. These negative results could be due to insufficient measurement of this dimension, or due to sexual harassment and unfair treatment being virtually absent in our samples. Indeed, 123 of the 348 external phase respondents opted not to respond to the sexual harassment items, which could be because they did not have these experiences or they were not comfortable reporting them. We recommend collecting larger or more targeted samples when focusing on these scales due to their low incidence.

## DISCUSSION

Prior research indicates that quality experiences with research mentors help to maximize the benefits of undergraduate research experiences (Aikens et al., 2016, 2017; Gentile et al., 2017; Hernandez et al., 2018; Joshi et al., 2019). However, recent research suggests that not all undergraduate research experiences are positive (Cooper et al., 2019; Gin et al., 2021) and negative experiences with research mentors may be an influential factor in the quality of undergraduates’ research experiences (Limeri, Asif, Bridges, et al., 2019). In light of national calls to broaden participation in undergraduate research, there is urgent need to determine the range of mentoring experiences by undergraduate researchers and to systematically investigate *who* experiences supportive and destructive mentoring and *how* these experiences influence students’ personal and professional growth. Through the work we describe here, we have produced a measure in accordance with best practices for establishing the validity and reliability of new measures (AERA et al., 2014), which will make this systematic investigation possible.

The MURS adds value beyond existing measures of mentoring, such as the Mentoring Competency Assessment (MCA) and Mentoring Relationship Quality (MRQ) measures examined here. The subscales of the MURS allow for examination of different types of negative mentoring experiences, equipping the community with a tool that can be used to address more nuanced questions about which mentoring experiences lead to which student outcomes. The MURS is also inclusive of negative mentoring experiences, not just positive. The MURS has diverse forms of validity evidence, including evidence that students generally interpret items on the MURS as expected and responses on the MURS relate to students’ integration into the scientific community. Finally, the MURS expands the toolbox scholars and practitioners can use to examine the interactions between mentor competence, mentee experiences, mentor-mentee relationships, and mentor and mentee outcomes.

Our results are encouraging because they indicate that most undergraduate researchers experience high levels of supportive mentoring and low levels of destructive mentoring. Only a small proportion of external phase respondents agreed or strongly agreed that they had mentors who were inaccessible or a mismatch, or who did not provide sufficient career and technical support. Destructive forms of mentoring, such as sexual harassment and unfair treatment, were observed even more rarely. That said, our results are also discouraging. Undergraduates in both of our samples reported experiencing the absence of supportive mentoring as well as destructive mentoring, and these negative experiences were associated with less favorable outcomes of participating in research. Substantial time and resources are invested in undergraduate STEM research experiences (Eagan et al., 2013; Gentile et al., 2017). Yet, negative mentoring appears to be limiting students’ growth, which may be driving away talent and ultimately undermining these investments.

Our results suggest that negative mentoring experiences reported on the MURS are not simply an artifact of students’ personality traits. This result was replicable across our datasets, which is noteworthy because it contradicts results from studies of mentoring in workplace settings that find associations between mentee personality and provision of mentoring support (Turban & Dougherty, 1994; Turban & Lee, 2007). Furthermore, this result is important because mentees who report negative mentoring may be criticized as being “too sensitive” or otherwise at fault for their negative mentoring.

Our results indicated that undergraduates’ attachment styles are either unrelated or weakly related to their negative mentoring experiences. It is worth noting that undergraduates with anxious attachment styles reported modestly more negative mentoring experiences. This result is consistent with prior research indicating that individuals with these forms of attachment report smaller and less satisfying social networks (Anders & Tucker, 2000). It may be that undergraduates who have anxious attachment styles may benefit from the provision of greater mentoring support, which mentors can cultivate through professional development (Pfund et al., 2015). Such students may also benefit from completing professional development targeted at building interpersonal communication skills, such as *Entering Research* or mentoring up (Branchaw et al., 2020; Lee et al., 2015).

We examined whether the mentor’s position influenced negative mentoring and found no association between undergraduates’ experiences of negative mentoring and mentor position type (faculty vs. non-faculty). Additional research should be done to determine if this result is replicable. If so, this suggests that having mentors with more mentoring experience, like faculty members, or a closer career stage, like graduate students or postdoctoral associates, does not result in fewer negative mentoring experiences for undergraduate researchers. This also implies that mentors at all levels could benefit from completing mentoring professional development in order to develop the skills necessary to mentor effectively and inclusively (Byars-Winston et al., 2018; Pfund et al., 2015; Womack et al., 2020). Future research could explore whether mentors differ in other ways that are influential for students.

Our results provide empirical evidence that the absence of supportive mentoring experiences and destructive mentoring experiences are related yet distinct, at least in undergraduate research settings. It is noteworthy that interpersonal mismatch loaded onto both of these higher-order factors, with a slightly higher loading onto the destructive factor. The higher loading may reflect the fact that the *interpersonal mismatch* items are negatively worded. The cross-loading may reflect the fact that mismatch is inherently dyadic, while all other forms of mentoring undergraduates reported were perceptions of mentor characteristics or behaviors (or lack thereof). This interpretation aligns with findings from research in corporate settings, which indicates that mentees experience two forms of negative mentoring: poor mentor behavior and poor dyadic fit (Eby & Allen, 2002b). Our model fit statistics indicated that treating interpersonal mismatch as related to both higher-order factors was superior to treating it as a separate factor. It may be that negative mentoring experiences reflect a developmental process in which mentees gauge fit or mismatch with their mentor if they experience abusive supervision or an absence of career or psychosocial support. In contrast, mentees who experience supportive mentoring and effective supervision then feel like they are well-matched with their mentor. Longitudinal research that examines these dimensions of negative mentoring over the duration of a research experience would be necessary to test these hypotheses.

Related to the notion of mismatch, recent research has shown that undergraduate STEM mentees’ psychological similarity with their mentors – meaning the perception of shared perspectives, values, and work habits (Turban & Jones, 1988) – relates to higher levels of psychosocial support, relationship quality, and commitment to STEM careers (Hernandez et al., 2017a, 2017b; Pedersen et al., 2022). It may be that interpersonal mismatch and psychosocial similarity are two ends of the same continuum (i.e., one construct) or two distinct constructs. For instance, mentees may feel they are similar to their mentors (or not), without feeling mismatched, or they may perceive the absence of similarity as an indicator of mismatch. Given the positive effects of psychological similarity reported elsewhere and the negative effects of mismatch observed here, future research should examine how these constructs relate as well as how they function to influence undergraduate researchers’ professional growth and research career pursuits. For instance, research has shown that a “birds of a feather” intervention (Gehlbach et al., 2016; Robinson et al., 2019), which highlights a dyad’s shared interests, can promote psychological similarity and relationship quality between undergraduate STEM mentees and their mentors (Hernandez et al., 2020, 2023). Such an intervention may set undergraduate researchers on a path toward developing quality relationships with research mentors and buffer against the perception of interpersonal mismatch.

In our view, the MURS offers multiple benefits as a measurement tool. First, the MURS is the only mentoring measurement tool with robust validity evidence, especially related to predicting the effects on students of experiencing negative mentoring. Second, our results indicate that the MURS can be used in its entirety to measure negative mentoring experiences collectively or by dimension or higher-order factor to gain insight about specific forms of supportive and destructive mentoring. For instance, using the abusive supervision scale alone or the collection of destructive mentoring experience scales could reveal the distinct effects of what are likely to be the most detrimental forms of negative mentoring. Using the two supportive experience scales (career and technical and psychosocial) may be useful to mentors in identifying areas to improve or in supporting mentees in asking for the particular forms of support they need (Lee et al., 2015). Given that we did not study the use of subscales in this way, it would be important to continue to collect validity evidence related to these uses. It may be that undergraduate researchers will respond differently to the scales in isolation. For instance, undergraduate researchers may be more hesitant to report abusive supervision, sexual harassment, or unfair treatment if these scales are presented outside of the context of the positively framed scales.

The dimensionality of the MURS also affords an opportunity to gain mechanistic insights about the influence of specific forms of negative mentoring on undergraduate researchers’ career motivations and decisions. For instance, we observed the strongest negative relationships between insufficient career and technical and psychosocial support and undergraduates’ positive emotions about research. These forms of support may be necessary for undergraduates to experience pride, accomplishment, or pleasure in their research, which in turn might prompt them to continue in research. In contrast, we observed the strongest positive relationships between abusive supervision and undergraduates’ negative emotions about research, suggesting that abusive mentor behavior may be a source of worry or stress that ultimately drives undergraduates out of research paths. Testing these hypotheses would require examining students’ emotions, intentions, and mentoring experiences over time.

Longitudinal research using the MURS would also be useful for examining the influence of mentoring on students’ integration into the scientific community as a recursive process. Our results indicate that undergraduates do not differ in their experiences with negative mentoring based on their scientific self-efficacy *before* they begin research. However, lower pre-research levels of scientific identity, research beliefs, and intentions were significantly and positively related to experiencing negative mentoring. As noted above, mentors may be responding differently to students who feel less integrated into the scientific community at the outset of their research experience. Alternatively, students who feel less integrated may be securing research experiences with less suitable mentors, leading to more negative experiences, or attending to evidence that research is the wrong path for them (i.e., poor experiences with mentors).

Finally, the MURS could be used to examine the antecedents and consequences of negative mentoring in undergraduate research. For instance, research in corporate settings suggests that formal or assigned mentoring relationships are less effective than informally developments relationships (Eby et al., 2013; Hernandez et al., 2017a). This raises the question of whether the process through which undergraduate research mentoring relationships are established (e.g., did they choose or were they assigned to the research and/or research mentor?) may influence whether undergraduates experience negative mentoring. Future research could also assess the effects of interventions aimed at fostering positive mentoring relationships, including mentoring professional development for mentors and mentoring-up development for mentees (Lee et al., 2015; Pfund et al., 2015). Finally, the MURS could be used by institutions to monitor the quality of undergraduate research mentoring occurring on campus and to evaluate the effectiveness of efforts to improve undergraduate research mentorship over time.

## Acknowledgements

We are grateful to Angela Byars-Winston, Cheryl Dickter, Lillian Eby, Paul Hernandez, and Justin Lavner for their guidance, advice, and feedback. We thank David Esparza, Levon Esters, Matthew Holtz, Laura Lunsford, Christine Pfund, Zachary Wood, and Michelle Ziadie for providing additional review of our items. We are also grateful to all of our institutional partners who helped us recruit participants, our participants for sharing their time and experiences, and Social Psychology of Research Experiences and Education group members for feedback on drafts of this manuscript. This work was supported in part by National Science Foundation Improving Undergraduate STEM Education Grant 1841061 and the Georgia Athletic Association Professorship for Innovative Science Education. Any opinions, findings, conclusions, or recommendations expressed in this material are those of the authors and do not necessarily reflect the views of any of the funding organizations.

## Footnotes

1 VSS is an index that assesses the degree to which the loading pattern reflects simple structure (items have a high loading on a single factor, and near-zero loadings on all other factors); factor solutions with simple structure are preferred; Velicer’s MAP aims to find the solution that minimizes the average residual covariances after systematic factor variance is controlled for; Empirical BIC assesses the likelihood of the model given the data controlling for the number of parameters in the model based on the solution’s χ^2^ and *df*, and – all else equal – prefers simpler models over more complex ones; the Sample Sized Adjusted BIC is similar but is based on the model log-likelihood rather than the χ2 and the number of parameters rather than the *df*; in addition, it is adjusted for any differences in sample size (which was not an issue here).

2 Although we make use of the broad category of “Asian,” we recognize that students who identify as Asian have a spectrum of experiences and more careful disaggregation by specific cultural or national identity is needed to understand these experiences. We make use of the broad category of “minoritized” to include students who identify as American Indian/Alaskan Native, African American or Black, Native Hawaiian/Pacific Islander, and Hispanic/Latine. Again, we recognize there are important differences between these groups and students have a range of experiences within and across racial and ethnic groups. Our intention with using these broad categories is explore whether there are any patterns shared across these groups.

## Notes

### Competing Interest Statement

The authors have declared no competing interest.

