## Supplemental material for "Development and Validation of the Mentoring in Undergraduate Research Survey"

**Supplemental Materials for:**
**Development and Initial Validation of the Mentoring in Undergraduate Research Survey**

Lisa B. Limeri, Nathan T. Carter, Riley A. Hess, Trevor T. Tuma, Isabelle Koscik, Alexander J. Morrison, Briana Outlaw, Kathren Sage Royston, Benjamin H. T. Bridges, Erin L. Dolan

Table of Contents


| Contents | Pages |
|---|---|
| Cognitive Interview Screening Survey | S2-S3 |
| Cognitive Interview Script and Questions | S4 |
| Structural Phase Survey | S5-S7 |
| Figure S1. Scree plot from structural phase analyses | S8 |
| External Phase Pre-survey items | S9-S11 |
| External Phase Post-survey items | S12-S15 |
| References | S16 |

**Cognitive Interview Screening Survey**

Thank you for your interest in our research! Before you respond to any questions, please take a minute to think about *your most recent research experience*.

How would you rate the overall quality of this research experience?

1. Mostly positive
2. Mostly negative
3. Some positive and some negative

*Now think about the person who was your research mentor during this experience.* This person could be a faculty member, graduate student, or someone else in the research group.

How would you rate your experience working with this person?

1. Mostly positive
2. Mostly negative
3. Some positive and some negative

We may want to contact you for an interview to help us refine a survey we are developing about undergraduate researchers' experiences with their research mentors, especially when those experiences are not so positive. If you choose to participate, you will receive a $25 gift card.

If you are willing, please enter your name and email address below. Please note that we will keep your information confidential but we need your name and contact information in order to schedule an interview.

First and last name:
Email Address:

The following questions are about your personal characteristics and will help the researchers ensure they have received input from students who represent a broad range of personal characteristics.

- What gender do you most closely identify with?
    - Female
    - Male
    - Transgender
    - Non-binary
    - Other: Please explain.
    - Prefer not to respond

- With which race(s) or ethnicity/ies do you most closely identify? Please choose all that apply.
    - American Indian, Alaskan Native, or Native American
    - Black or African American
    - East Asian (e.g., China, Japan, Korea)
    - Hispanic or Latino/Latina
    - Native Hawaiian or other Pacific Islander
    - North African or Middle Eastern
    - South Asian (e.g., India, Sri Lanka, etc)
    - White
    - Other: Please explain.
    - Prefer not to respond

- What type of research did you complete? Please choose all that apply.
  - Bench/wet
  - Field
  - Computational
  - Theoretical
  - Other: Please explain.
  - Prefer not to respond

- Please enter the full name of the college or university where you had the research experience you were responding about: (open response)

This information is confidential and will only be used to help the researchers make sure they collect responses from students who did research at different types of colleges and universities.

Thank you for completing this survey!

**Cognitive Interview Script and Questions**

Thanks for responding to our survey about the mentoring you experienced during your undergraduate research. You agreed to participate in this study when you responded to the survey, but I want to check in and see if you have any questions before we proceed. If you are comfortable, I'd like to record our conversation so I can be sure to capture your comments accurately and completely. Your comments will not be transcribed, and we will only use the audio recording in our analyses, so no one else will hear your voice or know that you participated in the study. We will also be taking notes to make sure we fully capture your feedback.

Are you comfortable with me recording our conversation?

- If yes, start recorder
- If no, then explain that we will be taking notes to make sure that we capture their comments as well as possible

Do you have any questions for me before we begin? [If no, proceed with interview.]

I'd like you to read each statement and respond from 1 strongly disagree to 5 strongly agree.
As you read each item, I'll ask you why you chose the rating you did and I might ask you to provide an example. I'll also ask you if there is anything about the statement that is confusing and whether you would change anything about the statement to make it clearer. Do you have any questions before we proceed?

Questions to be asked after the participant reads and responds to each item:

- Why did you choose that response?
- What does this statement mean to you? Can you give me an example?
- Is there anything about the statement that is confusing?
- Would you change anything about the statement to make it clearer?

Response scale:

1 Strongly Disagree
2 Disagree
3 Neither Agree nor Disagree
4 Agree
5 Strongly Agree

**Structural Phase Survey Items**

Please select your college or university [drop-down list]

**Mentoring in Undergraduate Research Survey draft items**
Prompt: Please indicate the extent to which you agree or disagree with each of these statements.
Response scale: 1 = "Strongly disagree"; 2 = "Somewhat disagree"; 3 = "Neither agree nor disagree"; 4 = "Somewhat agree"; 5 = "Strongly agree"

1. My mentor explained how my work fit into the bigger picture.
2. My mentor thought the work I did was important.
3. My mentor helped me understand the purpose of research tasks.
4. My mentor gave me work that was the right level of difficulty for me.
5. My mentor was friendly.
6. My mentor respected me.
7. My mentor was clear about how my performance was being evaluated.
8. My mentor gave me the attention I needed.
9. My mentor had faith in me.
10. My mentor valued my contributions to the research.
11. My mentor gave me the right amount of work.
12. My mentor cared about me as a person.
13. My mentor made sure I was prepared to do research tasks.
14. My mentor was clear about when I was expected to be working.
15. My mentor encouraged me.
16. My mentor gave me enough guidance in my research.
17. My mentor expected me to work reasonable hours.
18. My mentor and I talked about my career aspirations.
19. My mentor was available when I needed them.
20. My mentor was around to answer questions.
21. My mentor made time to meet with me.
22. My mentor responded when I contacted them.
23. My mentor gave me useful feedback on my work.
24. My mentor was clear about what they wanted me to do.
25. My mentor and I had incompatible work styles.
26. My mentor and I had a tense relationship.
27. My mentor created an intimidating environment.
28. My mentor was biased against certain groups of people.
29. My mentor and I had incompatible personalities.
30. My mentor was too harsh with their criticism.
31. My mentor and I worked poorly together.
32. My mentor discussed topics that were too personal.
33. My mentor was passive aggressive.
34. My mentor invaded my privacy.
35. My mentor made me do excessive grunt work.
36. My mentor made inappropriate comments about my personal life.
37. My mentor was condescending.
38. My mentor took credit for my work.
39. My mentor blamed me for their mistakes.
40. My mentor and I had difficulty getting along.
41. My mentor and I had incompatible communication styles.
42. My mentor had favorites in the lab.
43. My mentor treated people unfairly based on their... - Major

44. My mentor treated people unfairly based on their... - Race/ Ethnicity
45. My mentor treated people unfairly based on their... - Gender/ Sex
46. My mentor treated people unfairly based on their... - Religion
47. My mentor treated people unfairly based on their... - Sexual orientation
48. My mentor treated people unfairly based on their... - Career interests

Prompt: Please indicate the frequency of each behavior.
Response scale: 1 = "Never"; 2 = "Less than monthly"; 3 = "Weekly"; 4 = "Daily"
1. My mentor was rude to me.
2. My mentor gossiped about people in the lab.
3. My mentor contacted me about their personal problems.
4. My mentor belittled me.
5. My mentor scolded people in the lab.

Content warning: The next set of statements contains information about sexual harassment or misconduct, which may be triggering to survivors. You may choose to skip the next section. Support is available 24 hours a day at Project Safe Hotline: 615-322-SAFE(7233)
6. My mentor touched me without my permission.
7. My mentor made sexual remarks.
8. My mentor made sexual comments about me.
9. My mentor made sexual jokes.

**Personality**
Prompt: Please indicate how much the following statements describe you and your personality. Response scale: 1 = Very Inaccurate; 2 = Moderate Inaccurate; 3 = Neither Inaccurate nor Accurate; 4 = Moderately Accurate; 5 = Very Accurate; 6 = Prefer Not to Respond
Source & Evidence of validity: We used the mini-IPIP measure developed by Donnellan and colleagues (2006). In their paper, they report evidence of validity.
1. I am the life of the party.
2. I socialize with a lot of different people at parties.
3. I dominate conversations.
4. I keep in the background.
5. I sympathize with others' feelings.
6. I feel others' emotions.
7. I am not really interested in others.
8. I am not interested in other people's problems.
9. I get chores done right away.
10. I like order.
11. I often forget to put things back in their proper place.
12. I make a mess of things.
13. I have frequent mood swings.
14. I get upset easily.
15. I am relaxed most of the time.
16. I seldom feel blue.
17. I have a vivid imagination.
18. I have difficulty understanding abstract ideas.
19. I am not interested in abstract ideas.
20. I do not have a good imagination.

In what discipline did you conduct research?

- Response options: Life Sciences (e.g., ecology, biochemistry); Chemistry, Engineering and Computer Science; Physics, Geosciences, other, prefer not to respond

How much research experience have you had?

- Response options: None; One term; Two terms; Three terms; More than three terms; Prefer not to respond

What is your gender identity?

- Response options: Woman; Man; Another gender identity, please specify [write-in]; Prefer not to respond

What is your age in years? [write-in]

Please indicate the race(s) and ethnicity/ies with which you most closely identify. Select all that apply.

- Response Options: American Indian or Alaskan Native; Black or African American; East Asian; Hispanic or Latinx; Middle Eastern or North African; Native Hawaiian or Pacific Islander; South Asian; White; Other, please specify [write-in]; Prefer not to respond

Please indicate if you have any of the following impairments or disabilities (select all that apply).

- Response options: A sensory impairment (vision or hearing); A mobility impairment; A learning disability (e.g., ADHD, dyslexia); A mood disorder (e.g., depression, anxiety, bipolar); Autism Spectrum Disorder; A disability or impairment not listed above [write-in]; None of these; Prefer not to respond

What were your SAT Scores (if applicable): - Reading & Writing - Score
What were your SAT Scores (if applicable): - Math - Score
What were your ACT Scores (if applicable): - Composite Score - Score
Did you take the SAT before March of 2016?

- Response options Yes; No

What is the highest level of education that your parent(s)/guardian(s) have completed?

- Parent 1? Parent 2?
- Response options: did not complete high school; high school diploma or GED; some college; associates degree; bachelor's degree; master's degree; doctoral or professional degree; I don't know; not applicable

**Figure S1. Scree plot from structural phase analyses.**

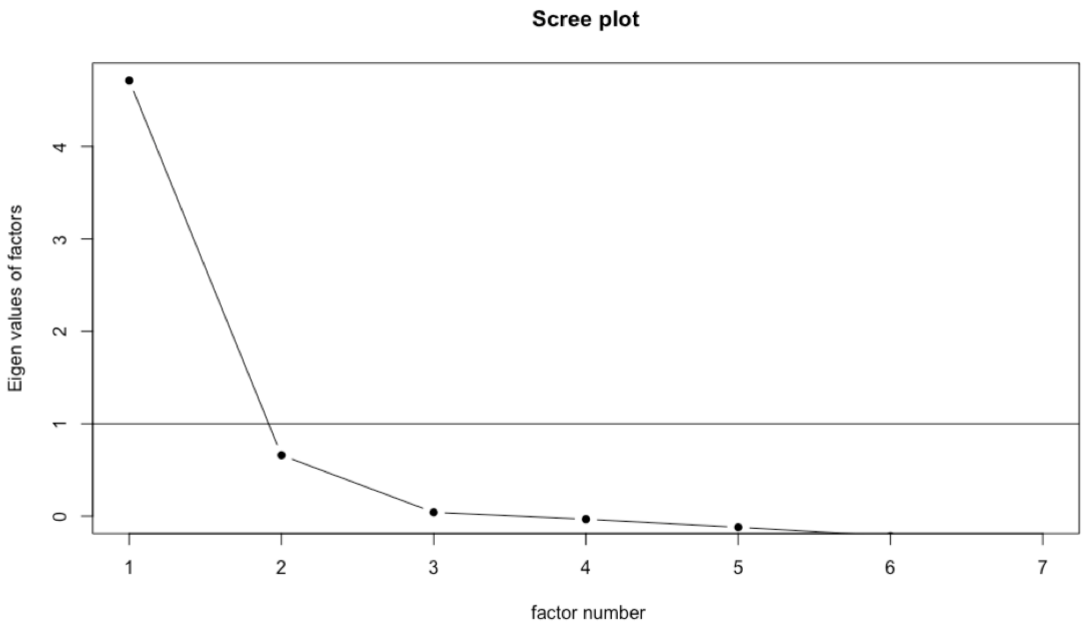

**External Phase Pre-survey Items**

Please select your college or university [drop-down list]

If you will be conducting research at an institution other than the one that you attend, please enter the institution name below. [write-in]

***Science Self-Efficacy***
Prompt: Please rate your confidence in your ability to perform the following tasks.
Response scale: 1 = Not At All Confident; 2 = Not Very Confident; 3 = Unsure; 4 = Fairly Confident; 5 = Absolutely Confident; 6 = Prefer Not to Respond
Source & Evidence of validity: This measure was developed by Chemers et al. (2011) and refined by Estrada et al. (2011). Both papers present evidence of validity based on internal structure and reliability.

1. Use technical science skills (use of tools, instruments, and/or techniques).
2. Generate a research question to answer.
3. Figure out what data/observations to collect and how to collect them.
4. Create explanations for the results of the study.
5. Use scientific literature and/or reports to guide research.
6. Develop theories (integrate and coordinate results from multiple studies).

***Science Identity***
Prompt: Please rate the extent to which you agree or disagree with the following statements.
Response scale: 1 = Strongly Disagree; 2 = Somewhat Disagree; 3 = Neither Agree nor Disagree; 4 = Somewhat Agree; 5 = Strongly Agree; 6 = Prefer Not to Respond
Source & Evidence of validity: This measure was developed by Chemers et al. (2011) and refined by Estrada et al. (2011). Both papers present evidence of validity based on internal structure and reliability.

1. I have a strong sense of belonging in the community of scientists.
2. I derive great personal satisfaction from working on a team that is doing important research.
3. I have come to think of myself as a scientist.
4. I feel like I belong in the field of science.
5. The daily work of a scientist is appealing to me.

***Value Beliefs about Research***
Prompt: Please rate the extent to which you agree or disagree with the following statements.
Response scale: 1 = Strongly Disagree; 2 = Somewhat Disagree; 3 = Neither Agree nor Disagree; 4 = Somewhat Agree; 5 = Strongly Agree; 6 = Prefer Not to Respond
Source & Evidence of validity: This measure was developed by Gaspard and colleagues (2015). Items were originally intended to measure students' values regarding math. The word "math" was replaced with "research." Gaspard et al., 2015 report evidence of validity based on internal structure.

<u>Intrinsic value</u>
1. Research is fun for me.
2. I like doing research.
3. I simply like research.
4. I enjoy dealing with research topics.

<u>*Communal value*</u>
1. Being well-versed in research will prepare me to help my community.
2. I can do good in the world based on my ability to do research.
3. If I can do research well, I can make a difference in the world.

<u>*Opportunity cost*</u>
1. I have to give up other activities that I like to be successful at research.
2. I have to give up a lot to do well in research.
3. I'd have to sacrifice a lot of free time to be good at research.

***Intentions***
Prompt: Rate the extent to which you are likely to pursue …
Response scale: 1 = Not at all likely; 2 = Somewhat Unlikely; 3 = Not Sure; 4 = Somewhat Likely; 5 = Very Likely; 6 = Prefer Not to Respond
Source & Evidence of validity: We measure intentions to pursue further education and a science and/or research career as an indicator of actual pursuit based on evidence that intent is a strong predictor of actual behavior (Eagan et al., 2013; Estrada et al., 2011). We measure intent to persist in science using three items adapted from Estrada and colleagues (2011), who report strong evidence that responses to these items correlate well with longitudinal behavioral indicators persistence.
1. A career in science.
2. A career in research.
3. Graduate education in science.

***Personality***
Prompt: Please rate how accurate each of these statements is for you.
Response scale: 1 = Very Inaccurate; 2 = Moderate Inaccurate; 3 = Neither Inaccurate nor Accurate; 4 = Moderately Accurate; 5 = Very Accurate; 6 = Prefer Not to Respond
Source & Evidence of validity: We used the mini-IPIP measure developed by Donnellan and colleagues (2006). In their paper, they report evidence of validity.

<u>*Neuroticism*</u>
1. I experience my emotions intensely.
2. I seldom get emotional.
3. I am not easily affected by my emotions.
4. I experience very few emotional highs and lows.

<u>*Openness*</u>
1. I am not interested in abstract ideas.
2. I avoid philosophical discussions.
3. I have difficulty understanding abstract ideas.
4. I am not interested in theoretical discussions.

***Attachment***
Prompt: Please rate the extent to which you agree or disagree with the following statements.
Response scale: 1 = Strongly Disagree; 2 = Somewhat Disagree; 3 = Neither Agree nor Disagree; 4 = Somewhat Agree; 5 = Strongly Agree; 6 = Prefer Not to Respond
Source & Evidence of validity: This Measure of Attachment Qualities (MAQ) was developed by Carver (1997). In a 25-year review of attachment measures, Ravitz et al. (2010) noted that the MAQ demonstrates strong validity and reliability evidence. Throughout, we replaced the words "my partner" with the word "others."

_Avoidant_
1. I get uncomfortable when someone wants to be very close.
2. I find it easy to be close to others.
3. I prefer not to be too close to others.
4. I am very comfortable being close to others.
5. Others want me to be more intimate than I feel comfortable being.

_Anxious_
1. I often worry that others don't really love me.
2. I often worry that others will not want to stay with me.
3. I don't worry about others abandoning me.
4. I have trouble getting others to be as close as I want them to be.
5. I find others often are reluctant to get as close as I would like.
6. My desire to merge sometimes scares people away.

_Secure_
1. When I'm close to someone it gives me a sense of comfort about life in general.
2. It feels relaxing and good to be close to someone.
3. Being close to someone gives me a source of strength for other activities.

**Sociodemographic questions**

In what discipline will you conduct research?
- Response options: Life Sciences (e.g., ecology, biochemistry); Chemistry; Engineering or Computer Science; Physics; Geosciences; Other: [write-in]; Prefer Not to Respond

What is your gender identity?
- Response options: Woman; Man; Another gender identity [write-in]; Prefer not to respond

What is your age in years? [write-in]

Please indicate the race(s) and ethnicity/ies with which you most closely identify. Select all that apply.
- Response options: American Indian or Alaskan Native; Black or African American; East Asian; Hispanic or Latinx; Middle Eastern or North African; Native Hawaiian or Pacific Islander; South Asian; White; Other, please specify [write-in]; Prefer Not to Respond

Please indicate if you have any of the following impairments or disabilities (select all that apply).
- Response options: A sensory impairment (vision or hearing); A mobility impairment; A learning disability (e.g., ADHD, dyslexia); A mood disorder (e.g., depression, anxiety, bipolar); Autism Spectrum Disorder; A disability or impairment not listed above; None of these; Prefer not to respond

What is the highest level of education that your parent(s)/guardian(s) have completed?
- Parent 1? Parent 2?
- Response options: did not complete high school; high school diploma or GED; some college; associates degree; bachelor's degree; master's degree; doctoral or professional degree; I don't know; not applicable

**External Phase Post-survey Items**

This survey will ask you to think about one person who mentored you in doing research. This could be the person you worked with most closely or the person who influenced you the most, either positively or negatively. What position did this mentor hold?

- Response options: “Faculty member. This is the principal investigator or head of the group in which you are conducting or have conducted research.”; “Postdoctoral associate. This is a member of the lab who has already completed a Ph.D.”; “Graduate student. This is a member of the lab who is earning a graduate degree.”; “Undergraduate student. This is a member of the lab who is earning an undergraduate degree.”; Other - please explain” [write-in]; “Prefer not to respond”

Please indicate the quality of mentoring you received from that person.

- Response options: Mostly positive; Some positive, some negative; Mostly negative; Prefer not to respond

***Mentoring in Undergraduate Research Survey*** *full draft item set*
Prompt: Please rate the extent to which you agree or disagree with the following statements.
Response Scale: 1 = "Strongly disagree"; 2 = "Somewhat disagree"; 3 = "Neither agree nor disagree"; 4 = "Somewhat agree"; 5 = "Strongly agree"
Source & Evidence of validity: These items were for the current study and evidence of validity is presented in the current manuscript.

*Abusive Supervision*

1. My mentor discussed topics that were too personal.*
2. My mentor was passive aggressive.
3. My mentor invaded my privacy.*
4. My mentor made me do excessive grunt work.
5. My mentor made inappropriate comments about my personal life.
6. My mentor was condescending.
7. My mentor took credit for my work. *
8. My mentor blamed me for their mistakes.
9. My mentor created an intimidating environment.
10. My mentor was too harsh with their criticism.
11. My mentor told me too much about their personal life.
12. My mentor was rude to me.
13. My mentor gossiped about people in the lab.*
14. My mentor belittled me.
15. My mentor scolded people in the lab.*

*Accessibility* (reverse scored)

1. My mentor was available when I needed them.
2. My mentor was around to answer questions.
3. My mentor made time to meet with me.
4. My mentor responded when I contacted them.
5. My mentor gave me the attention I needed.

*Big Picture Support* (reverse scored)

1. My mentor told me how the research in our group makes a difference.
2. My mentor explained how my work contributes to scientific knowledge.
3. My mentor made sure I knew that the work I was doing was scientifically valuable.
4. My mentor described how my project fit into the work of the research group.
5. My mentor helped me understand the purpose of my research tasks.
6. My mentor helped me see how my work fit into the bigger picture. *

*Career & Technical Support* (reverse scored)

1. My mentor made sure I was prepared to do research tasks.
2. My mentor gave me enough guidance in my research.
3. My mentor and I talked about my career aspirations. *
4. My mentor gave me useful feedback on my work.
5. My mentor gave me work that was the right level of difficulty for me.
6. My mentor was clear about how my performance was being evaluated.
7. My mentor gave me the right amount of work.
8. My mentor was clear about what they wanted me to do.
9. My mentor was clear about when I was expected to be working.*
10. My mentor expected me to work reasonable hours. *

*Psychosocial Support* (reverse scored)

1. My mentor thought the work I did was important. *
2. My mentor was friendly.
3. My mentor respected me.
4. My mentor had faith in me.
5. My mentor valued my contributions to the research.
6. My mentor cared about me as a person.
7. My mentor encouraged me.

*Interpersonal Mismatch*

1. My mentor and I had incompatible work styles. *
2. My mentor and I had a tense relationship.
3. My mentor and I had incompatible personalities.
4. My mentor and I worked poorly together.
5. My mentor and I had difficulty getting along.
6. My mentor and I had incompatible communication styles.

*Unfair Treatment*

1. My mentor was biased against certain groups of people. *
2. My mentor had favorites in the lab. *
3. My mentor treated people unfairly based on their... – Major*
4. My mentor treated people unfairly based on their... - Race/ Ethnicity
5. My mentor treated people unfairly based on their... - Gender/ Sex
6. My mentor treated people unfairly based on their... - Religion
7. My mentor treated people unfairly based on their... - Sexual orientation
8. My mentor treated people unfairly based on their... - Career interests*

* These items were excluded from the final item set.

**Mentoring Competency Assessment (MCA)**
Prompt: Please rate how skilled your mentor is in the following areas:
Response Scale: 1 = "Strongly disagree"; 2 = "Somewhat disagree"; 3 = "Neither agree nor disagree"; 4 = "Somewhat agree"; 5 = "Strongly agree"
Source & Evidence of validity: These items were developed by Fleming and colleagues (2013). Evidence for validity is presented in their original paper.

*Effective Communication*
1. Active listening.
2. Providing you with constructive feedback.
3. Establishing a relationship based on trust with you.
4. Identifying and accommodating different communication styles.
5. Employing strategies to improve communication with you.
6. Coordinating effectively with other mentors with whom you work.

*Aligning Expectations*
1. Working with you to set clear expectations of the mentoring relationship.
2. Aligning his/her expectations with your own.
3. Considering how personal and professional differences may impact expectations.
4. Working with you to set research goals.
5. Helping you develop strategies to meet research goals.
6. Accurately estimating your level of scientific knowledge.
7. Accurately estimating your ability to conduct research.
8. Employing strategies to enhance your understanding of research.

*Fostering Independence*
1. Motivating you.
2. Building your confidence.
3. Stimulating your creativity.
4. Acknowledging your professional contributions.
5. Negotiating a path to professional independence with you.

*Addressing Diversity*
1. Taking into account the biases and prejudices s/he brings to your mentor/mentee relationship.
2. Working effectively with mentees whose personal background is different from his/her own (age, race, gender, class, region, culture, religion, family composition, etc.)

*Promoting Professional Development*
1. Helping you network effectively.
2. Helping you set career goals.
3. Helping you balance work with your personal life.
4. Understanding their impact as a role model for you.
5. Helping you acquire resources (e.g., grants, etc.).

**Mentoring Relationship Quality (MRQ) Scale**
Prompt: Please rate the extent to which you agree or disagree with the following statements.
Response Scale: 1 = "Strongly disagree"; 2 = "Somewhat disagree"; 3 = "Neither agree nor disagree"; 4 = "Somewhat agree"; 5 = "Strongly agree"
Source & Evidence of validity: These items were developed by Allen and Eby (2003). Evidence for validity is presented in their original paper.

1. The mentoring relationship between my mentor and I was very effective.
2. I am very satisfied with the mentoring relationship my mentor and I developed.
3. I was effectively utilized as a mentee by my mentor.
4. My mentor and I enjoyed a high-quality relationship.
5. Both my mentor and I benefited from the mentoring relationship.

**Sexual Harassment** (Aycock et al., 2019)
Prompt: Please indicate the frequency of each behavior.
Response Scale: 1 = "Never"; 2 = "Less than monthly"; 3 = "Weekly"; 4 = "Daily"
Source & Evidence of validity: These items were developed by Aycock and colleagues (2019) in the context of physics. Evidence for validity is presented in their original paper. We modified the wording to reference a mentor specifically.

These items were also preceded by the following content warning and respondents were given the option to skip the items: The next set of statements contains information about sexual harassment or misconduct, which may be triggering to survivors. You may choose to skip the next section. Support is available 24 hours a day at Project Safe Hotline: 615-322-SAFE(7233)

1. My mentor touched me without my permission.
2. My mentor made sexual remarks.
3. My mentor made sexual comments about me. (excluded from the final item set)
4. My mentor made sexual jokes.

**Science Self Efficacy, Science Identity, Value Beliefs, and Intentions**
Same items as appeared on pre-survey

**Emotions about Research**
Prompt: How much do these words describe your **feelings about doing research?**
Response Scale: 1 = "Not at all"; 2 = "A little"; 3 = A moderate amount"; 4 = "A lot"; 5 = "A great deal"
Source & Evidence of validity: This measure was developed by Clark et al., (2016). They provide evidence of validity and sourced the emotions on the checklist from Larsen & Kasimatis (1990).

1. Proud
2. Stressed
3. Accomplished
4. Bored
5. Apathetic
6. Pleased
7. Worried
8. Enthusiastic
9. Annoyed
10. Excited
11. Exhausted
